# Post-Domestication selection of MKK3 Shaped Seed Dormancy and End-Use Traits in Barley

**DOI:** 10.1101/2025.07.11.664137

**Authors:** Morten E. Jørgensen, Dominique Vequaud, Yucheng Wang, Christian B. Andersen, Micha Bayer, Amanda Box, Katarzyna B. Braune, Yuanyang Cai, Fahu Chen, Jose A. Cuesta-Seijo, Haoran Dong, Geoffrey B. Fincher, Zoran Gojkovic, Zihao Huang, Benjamin Jaegle, Sandip M. Kale, Flavia Krsticevic, Pierre-Marie Le Roux, Antoine Lozier, Qiongxian Lu, Martin Mascher, Emiko Murozuka, Shingo Nakamura, Martin Ude Simmelsgaard, Pai R. Pedas, Pierre A. Pin, Kazuhiro Sato, Manuel Spannagl, Magnus W. Rasmussen, Joanne Russell, Miriam Schreiber, Hanne C. Thomsen, Nina W. Thomsen, Sophia Tulloch, Cynthia Voss, Birgitte Skadhauge, Nils Stein, Eske Willerslev, Robbie Waugh, Christoph Dockter

**Author notes:** Current address, Novo Nordisk A/S; 2880 Bagsværd, Denmark (Jose A. Cuesta-Seijo). Current address, Crop Genetics and Biotechnology Department of Agroecology, Aarhus University; Slagelse, Denmark (Sandip M. Kale). Current address, DLF Seeds A/S; 4000 Roskilde, Denmark. These authors contributed equally: Morten E. Jørgensen, Dominique Vequaud, Yucheng Wang.

## Abstract

Anthropogenic selection of grain traits such as dormancy has shaped the developmental trajectories of crop plants *(1)*. In cereals, shortening dormancy provides rapid and even post-harvest germination, but increases the risk of weather-induced pre-harvest sprouting (PHS) with harvest losses estimated at beyond 1 billion USD a year *(2, 3, 4)*. Our understanding of how, why, when and where diversification of cereal dormancy arose is fragmentary. Here, we show in the founder cereal crop barley *(Hordeum vulgare)* that the Mitogen-Activated Protein Kinase (MAPK) pathway regulates dormancy primarily through a mosaic of locus haplotypes comprising copy-number variation and inherent kinase activity of *Mitogen-activated protein kinase kinase 3 (MKK3)*. We provide evidence supporting the historical selection of specific *MKK3* haplotypes that shape dormancy levels according to changing climatic pressures. Understanding the regulatory landscape of MKK3 provides a genetic framework to balance short grain dormancy and PHS-avoidance during a period of rapid climate change.

## Main Text

Crops selected for enhanced overall performance, improved seed characteristics and adaption to a wide range of natural environments drove the emergence of agriculture and provided the foundation for advanced human societies (*1, 5*). Understanding how crop plants adapted to both anthropogenic and environmental demands over recent evolutionary time could therefore be central to developing long-term sustainable agricultural practices that cope with emerging global threats including our rapidly changing climate. However, the molecular mechanisms that underlie the successful adaptation of many key crop traits to different pressures remains poorly understood.

Dormancy in cereal grains (i.e. the suppression of germination in viable grains under environmental conditions normally favourable for germination) is one such trait. In wild cereal ancestors, adaptation to unpredictable or seasonal habitats has promoted variability in dormancy levels, a vital and evolutionary conserved bet-hedging strategy to ensure reproductive success of a population in case of catastrophic events (6). During domestication, anthropogenic selection shortened dormancy because grain with rapid and uniform germination optimised cultivation (*6*), while simplifying grain storage and subsequent processing. After spreading to temperate and sub-tropical areas, short dormancy allowed for direct resowing of freshly harvested grain, enabling the production of two crops per year and significantly increasing annual yield (*7-11*). However, short dormancy may also promote the undesirable outcome of uncontrolled germination (sprouting) of mature grain before harvest in response to warm and moist weather, a phenomenon called pre-harvest sprouting (PHS) (Fig. 1A). PHS (*2*) reduces the quality of mature grain and jeopardises its end-use value. Selection for short dormancy can thus have either beneficial or detrimental outcomes.

**Fig. 1.**
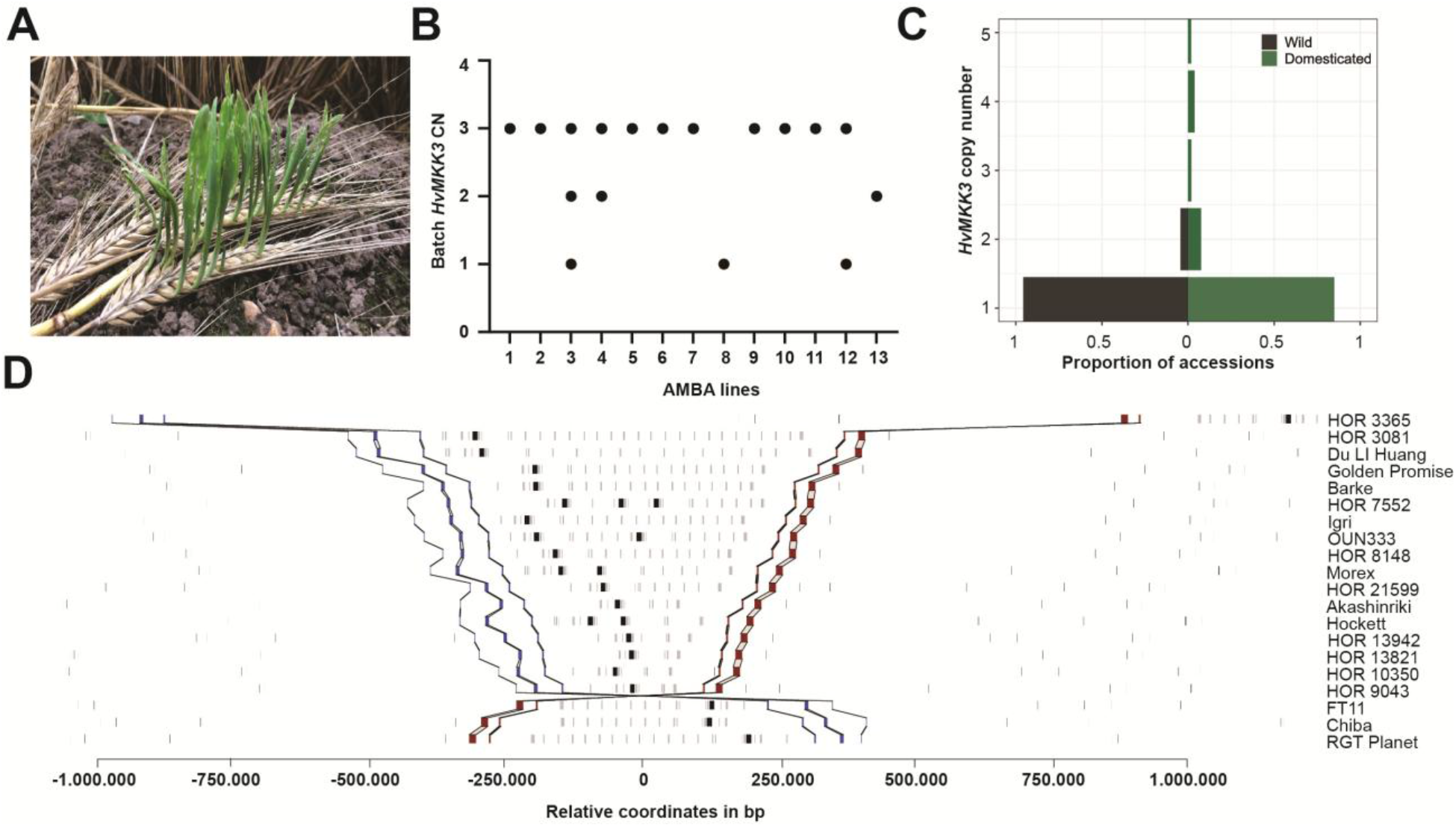
*MKK3* locus complexity in barley. **A**, Pre-harvest sprouted barley inflorescence. **B**, ddPCR *MKK3* copy number (CN) analysis across grain batches of AMBA-recommended malting barleys (Supplementary Table S1). **C**, Distribution of *MKK3* CN (as proportion of wild (23) or domesticated (53) accessions) across 76 pangenome assemblies. **D**, *MKK3* CNs and structural variations in 20 selected genome assemblies (add REFs BPGv2 and Pantranscriptome*9*). *MKK3* genes in black squares, other genes in grey squares. Red and blue squares denote marker genes that define the synteny, delimit the region and sort the accessions based on the distance between endpoints. Lines connect gene models between different genomes. Accession names are given on the right axis. In three accessions (FT11, Chiba, and RGT Planet), the larger *MKK3* region is inverted (6, 4 and 4 Mb, respectively).

During the development of modern agriculture, alleles causing shorter grain dormancy frequently became fixed to realise associated benefits. An unintended consequence is that environmentally triggered PHS continued to affect cereal crops in certain years and locations (*11-18*), downgrading cereal grain quality and value, with yearly losses in the billion-dollar range (*3, 4*). With a predicted rise in atmospheric temperature alongside an increased frequency of extreme weather events (*19*), the incidence of PHS and associated crop loss will likely increase (*20*), intensifying the current food systems crisis (*21*). How to balance the avoidance of PHS with the benefits of reduced dormancy in cereals is therefore a high priority.

In the founder cereal crop barley, dormancy is considered a quantitative trait with a major genetic component identified as *MKK3*, which exhibits diverse DNA sequence haplotypes reported to correlate with phenotypic variation (*12-14, 22*). By dissecting the molecular basis of *MKK3* haplotype variation in barley genome diversity panels combined with pre-breeding and multi-year field trial validation we provide new understanding of the directed evolution of cereal grain dormancy via environmental and cultural practices. We explore the origin and application of functionally diverse *MKK3* haplotypes, with the aim of implementing genome-enabled breeding for balanced and sustainable high-performance agriculture in changing environments (*23, 24*).

### Grain dormancy and PHS in barley

In barley, a single amino acid exchange in MKK3 (MKK3^T260^, i.e. Asparagine (N) to Threonine (T) at position 260), relative to the genomic reference HORVU.MOREX.PROJ.5HG00474740 (MKK3^Ref^), lowers MKK3 kinase activity *in vitro* (*13*). This variant is associated with increased grain dormancy and PHS avoidance in East Asian regions where wet harvests regularly promote PHS (*13*). In regions with a dry harvest climate such as the interior plains of North America, elite cultivars are associated with the opposite phenotype; low grain dormancy and high susceptibility to PHS in occasional wet harvest years. Genetic analyses (*15, 16, 25-27*) have shown that PHS susceptibility in this genetic material is correlated with an *MKK3* haplotype with a Glutamic Acid (E) to Glutamine (Q) exchange at position 165 (i.e. MKK3^Q165^) (*26, 27*). To explore the generality of this observation we investigated *MKK3* sequence variation in the PHS-susceptible North American landmark cultivars (*cvs.*) Klages, Harrington and CDC Meredith and confirmed the presence of MKK3^Q165^. However, the DNA sequencing signals appeared heterozygous for the mutation (Supplementary Fig. S1A). As barley cultivars are homozygous inbred lines, we therefore checked *MKK3* for copy number variation (CNV), a common source of heterozygous genotyping calls, using droplet digital PCR (ddPCR). We found that *MKK3* was triplicated in all three accessions, with the MKK3^Q165^-encoding variant present as two copies, thus explaining the heterozygous sequence signals (Supplementary Table S1). We then extended our CNV analysis to current American Malting Barley Association (AMBA) recommended varieties (*26*) (Supplementary Table S1). We found both increased *MKK3* copy number and the MKK3^Q165^-encoding variant in most AMBA lines and additionally CNV within batches of grain (Fig. 1B), an intriguing observation that merits further investigation. This indicates that the *MKK3* locus is more complex than previously reported and that heterogeneous *MKK3* haplotypes persist in elite breeding germplasm. Proposed genotype-phenotype associations for *MKK3* will therefore be inaccurate (*13, 25*), compromised by the genomic architecture of the *MKK3* locus and undermining breeding progress for PHS resilience (*27*).

### *MKK3* locus is complex in barley

To resolve these observations, we examined the *MKK3* locus in the barley pangenome (BPGv2) (*28*). Within the 76 pangenome sequences, we identified a total of 93 *MKK3* genes (defined as start to stop codon, including introns). Of these, 48 are distinct gene sequences. Restricting our analysis to coding sequences (CDS) revealed 24 CDS haplotypes encoding 13 unique MKK3 proteins (Supplementary Table S2). In 22 out of 23 (22/23) wild barley (*Hordeum spontaneum*) accessions, we found a single full-length *MKK3* gene copy, compared with up to 5 tandemly duplicated or structurally re-arranged *MKK3* copies in domesticated barley as multiple sequence haplotypes (Fig. 1C;Fig. 1D; Supplementary Table S2; Supplementary Fig. S2, S3A). Extending our analysis to a diverse collection of 365 ddPCR genotyped, 1,342 whole genome sequenced and 228 exome sequenced barley lines we found that wild barley accessions predominantly contain a single *MKK3*, while domesticated barleys show extensive CNV, with up to 15 copies including multi-copy mixed sequence haplotypes (Fig. 2A; Supplementary Tables 1, 3, 4).

**Fig. 2.**
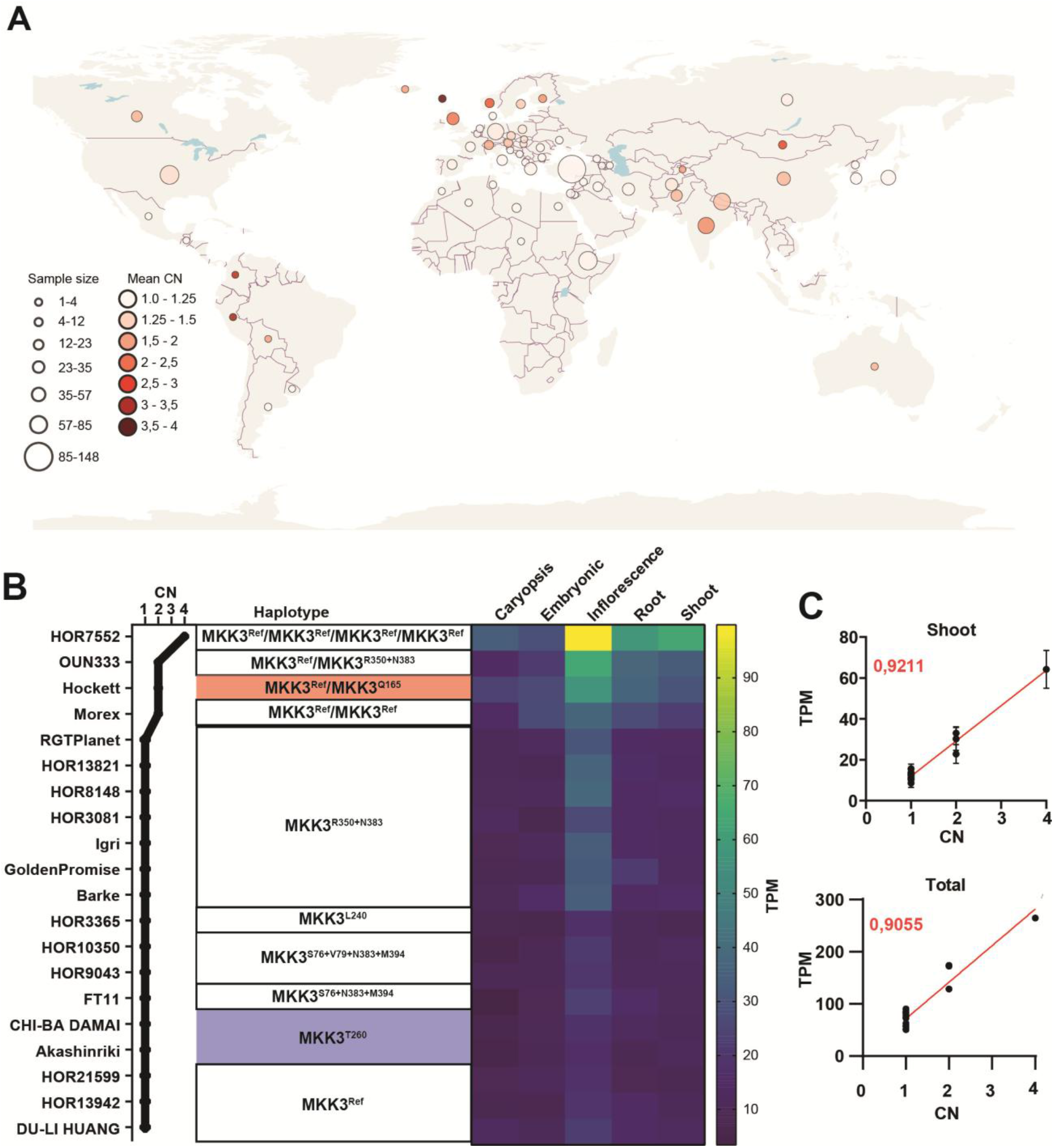
Global CN distribution of *MKK3* in barley and its functional impact. **A**, Mean *MKK3* CN across a global panel of 1,285 geo-localised barley accessions from 68 countries (420 accessions without recorded country of origin from Supplementary Table S3 are excluded). **B**, *MKK3* transcript abundance observed by mapping against the PanBaRT20 Reference Transcript Dataset (sum of chr5H56507 and chr5H56511) associated with *MKK3* copy number (CN). **C**, *MKK3* transcript abundance in the pantranscriptome (*29*) shoot tissue as a function of copy number with r^2^ value of the linear regression fit in red. Dot size shows sample size (in (**A**) ranging from 1-148 and in (**B**) 1-130 barley accessions.

We next explored the functional impact of the observed CNV by analysing transcript abundance variation in genotypes from the barley pan-transcriptome (*29*) (Fig. 2B; Supplementary Table S5). We found that firstly, European single-copy haplotypes (MKK3^R350+N383^, [i.e., Glycine (G) to Arginine (N) at position 350 and Aspartic Acid (D) to Asparagine (N) at position 383], e.g., *cv.* RGT Planet) show a tendency for slightly higher *MKK3* transcript abundance compared to accessions associated with high dormancy, such as wild barleys (e.g., B1K-04-12 [FT11]), some landraces (e.g., HOR13942) and East Asian barleys (e.g., Akashinriki with MKK3^T260^) (Fig. 2B; Supplementary Text 1.1). Secondly, MKK*3* copy number is positively correlated with transcript abundance (Fig 2C; Supplementary Fig. S3B). Thus, minor differences in transcript abundance are observed among genotypes containing a single *MKK3*, but fold changes are a direct consequence of CNV in domesticates.

We then examined the global MKK3 haplotype diversity in 1,071 geo-referenced domesticated barley accessions (Fig 3A; Supplementary Table S3), revealing distinct geographic patterns of haplotype enrichment. The high-dormancy variant MKK3^T260^ exhibited broader geographic distribution in regions with high precipitation and greater haplotype complexity than previously reported (Fig. 3A) (13,30). Notably, MKK3^T260^ was identified in combination with additional variants within both single-and multi-copy haplotypes (Supplementary Table S3), suggesting multiple independent evolutionary events, which potentially occurred as secondary adaptations to restore dormancy in domesticated barley as a consequence of range expansion. In contrast, the MKK3^Q165^ (low dormancy) multi-copy variant is predominantly found across Northern Europe and North America in 83 out of 1,707 accessions (Supplementary Tables 3; Fig. 2B), but also in landraces with a single *MKK3* (HOR 10775 in CORE1000; HOR 10892 in BPGv2) (28) (Fig. 3A). A median-joining haplotype network analysis of BPGv2 lines indicates that MKK3^Q165^ arose independently as single-and multi-copy locus haplotypes (Fig. 3B), suggesting that this variant causes a desired trait. The widely distributed MKK3^Ref^ is common across Europe, Asia and the Americas, whereas 41/43 MKK3^V79^ (i.e., Alanine (A) to Valine (V) at position 79) haplotypes are found in Ethiopian landraces. MKK3^N383^ and MKK3^R350^ variants are more common in Europe (Fig. 3A). Interestingly, the MKK3^N383^ variant is found in *Hordeum spontaneum* (wild barley) and in barleys secondary gene pool (*Hordeum bulbosum*) suggesting that the alternative allele MKK3^D383^ is a recently selected variant to increase dormancy (Supplementary Text 1.3) (31).

**Fig. 3.**
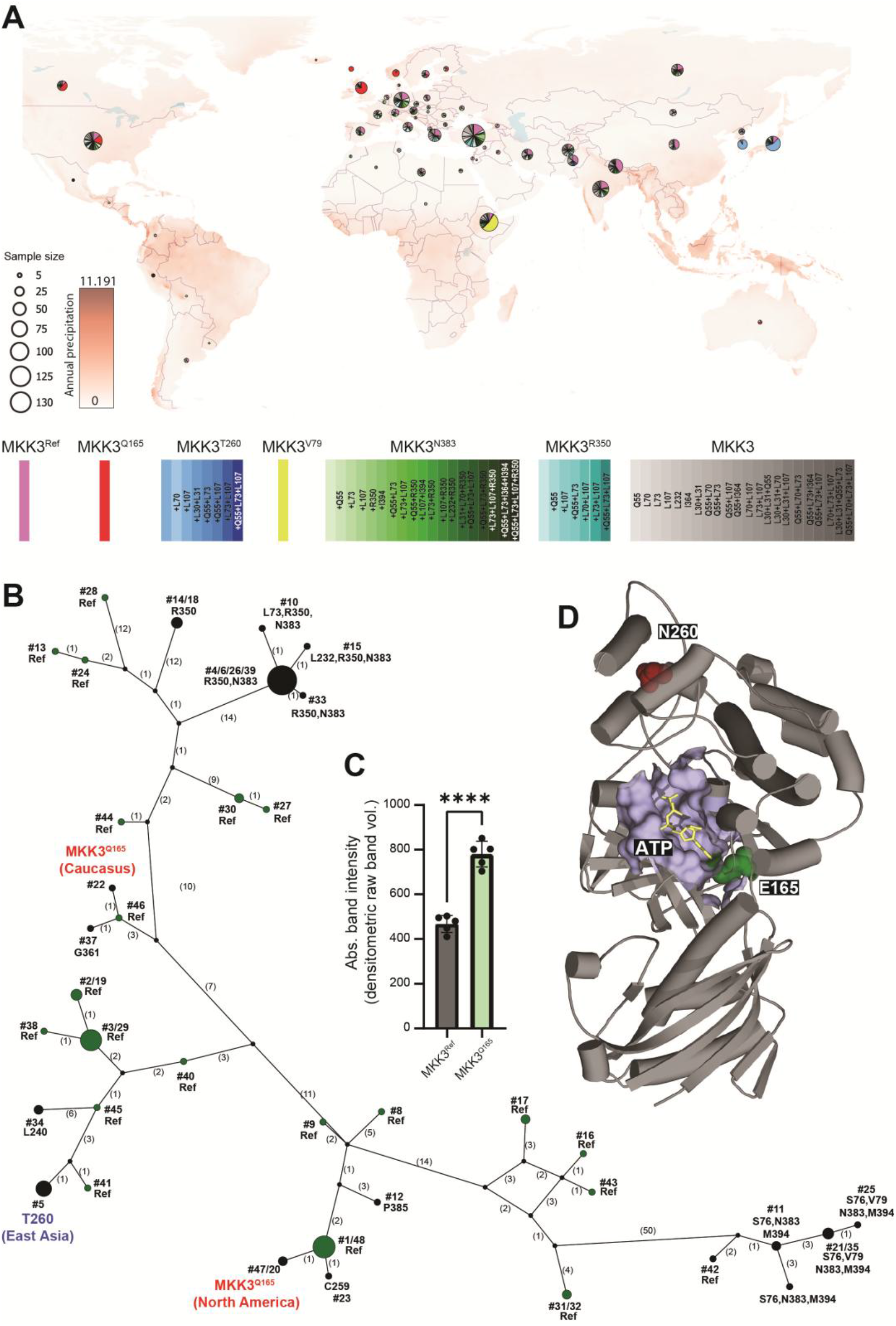
*MKK3* haplotype diversity and *in vitro* kinase activity. **A**, Global *MKK3* haplotype diversity across 1,071 geo-localized barley accessions (636 accessions without recorded country of origin, haplotype information or unique *MKK3* haplotypes (1 accession pr. haplotype) from Supplementary Table S3 were excluded). Annual mean precipitation (mm) (WorldClim2.1 BIO12(30) is shown as a red overlay gradient. Individual MKK3 haplotypes are color coded with MKK3^Ref^ in purple, MKK3^Q165^ in red, MKK3^T260^ containing in blue tones, MKK3^V79^ in yellow, MKK3^N383^ containing in green tones, MKK3^R350^ containing in teal tones and remaining MKK3 haplotypes as a grey tone. Dot size shows sample size. **B**, Median-joining haplotype network of *MKK3* copies in 76 pangenome assemblies. Node numbering represents different gene haplotypes (Supplementary Table S2). The node size is proportional to the number of gene IDs a given node represents. MKK3^Q165^ are shown in red, MKK3^T260^ are shown in blue. Nodes with MKK3^Ref^ are shown in green. Remaining MKK3 variants, compared to the amino acid haplotype #1 MKK3^Ref^ (Ref sequence, HORVU.MOREX.PROJ.5HG00474740) found in the extended barley pangenome in black (see haplotype numbering, Supplementary Table S2). **C,** *In vitro* kinase activity of MKK3^Ref^ (Supplementary Table S2) and variant MKK3^Q165^. Error bars are ±SD; P values (*P ≤ 0.05, ** P ≤ 0.01, *** P ≤ 0.001, and **** P ≤ 0.0001; ns, not statistically significant). **D**, MKK3 protein model (Alphafold & Alphafill model, based on reference MKK3^Ref^) showing amino acid position E165 (green spheres) and N260 (red spheres) associated with changes in MKK3 kinase activity. ATP binding site in kinase domain is shown in light blue with ATP (yellow sticks).

We next analysed the link between specific MKK3 amino acid variants and inherent kinase activity. Using *in vitro* kinase assays (see Methods) we confirm that MKK3^T260^ decreases kinase activity and show that MKK3^Q165^ increases kinase activity (Fig. 3C; Supplementary Fig. S3C). Molecular modelling leads us to propose that Q165 promotes the evolutionary conserved protein kinase DFG amino acid motif towards an active state, increasing kinase activity (*32*) (Fig. 3D; Supplementary Text 1.3). Testing the other common amino acid exchanges MKK3^N383^ and MKK3^V79^revealed that these show mild to strong increases in kinase activity, respectively (Supplementary Fig. S3D; Supplementary Text 1.3). We hypothesise that a combination of specific amino acid substitutions that alter MKK3 activity combined with CNV fine-tune overall MKK3 activity and consequently grain dormancy.

### Variation at *MKK3* affects dormancy and PHS susceptibility

To assess the practical impact of different *MKK3* locus haplotypes on dormancy and PHS we conducted field trials over several seasons with half of the plots of each genotype harvested at maturity in dry conditions and the other half exposed to rain and/or spray irrigation to promote the initiation of PHS (Fig. 4A). Harvested grain material was analysed using well-established tests from the barley malting industry; Germination Energy (GE) to measure grain dormancy and Germination Index (GI) to measure the speed of germination (see Methods). In 2023, we compared *cv.* RGT Planet (MKK3^R350+N383^), *cv.* Morex (MKK3^Ref^/MKK3^Ref^) and *cv.* Harrington (MKK3^Ref^/MKK3^Q165^/MKK3^Q165^) dry harvest samples, assaying GE and GI at 2 and 10 weeks after harvest. Dormancy was fully broken after 10 weeks for *cv.* RGT Planet, while *cvs.* Harrington and Morex, with increased *MKK3* copy number and with/without hyperactive MKK3^Q165^, respectively, had little grain dormancy (Fig. 4B).

**Fig. 4.**
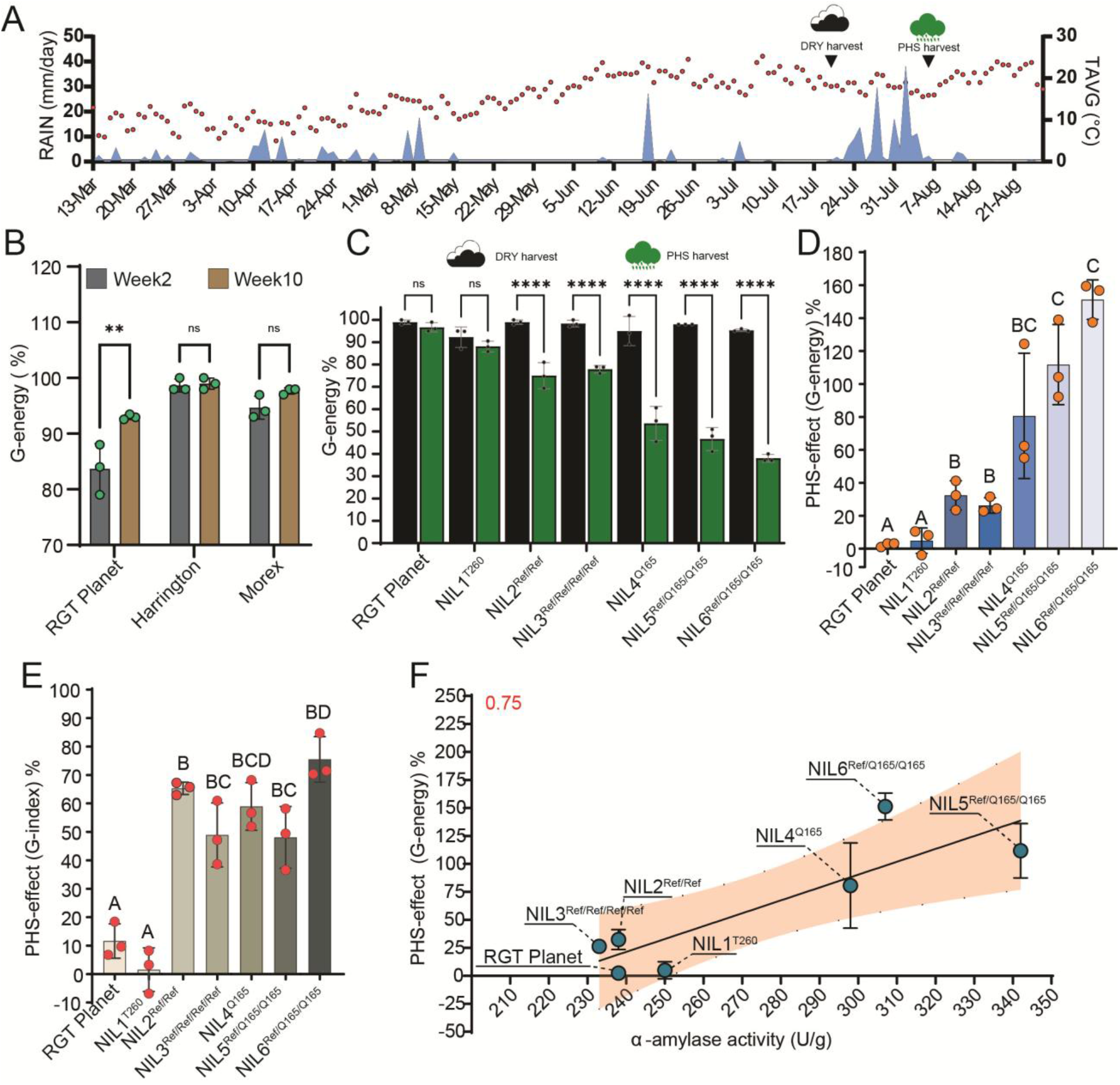
PHS field trials and analyses of diverse *MKK3* haplotype NILs. **A**, Weather data [the average air temperature (TAVC) in degree Celsius in red circles and precipitation (RAIN) in mm/day in blue shading] for PHS field trial site in FRA in the growth season 2023. Marked with arrows are the harvest time points of ‘dry’ and ‘PHS’ samples. **B**, G-energy of *cvs.* RGT Planet, Harrington and Morex at week 2 and 10. Error bars are ±SD; P values (*P ≤ 0.05, ** P ≤ 0.01, *** P ≤ 0.001, and **** P ≤ 0.0001; ns, not statistically significant). **C**, PHS trials of *cv.* RGT Planet (MKK3^R350+N383^), NIL1^T260^, NIL2^Ref/Ref^, NIL3 ^Ref/Ref/Ref/Ref^, NIL4 ^Q165^, NIL5 ^Ref/Q165/Q165^, NIL6 ^Ref/Q165/Q165^ (field-grown, FRA 2023). Shown are grain G-energy (percentage) after *cv.* RGT Planet dormancy is broken. Dry harvest (black bar) and wet harvest (green bar) samples. **D,E**, calculated PHS effect (G-energy) (**D**) and PHS effect (G-index) (**E**) of *cvs.* RGT Planet, Harrington and Morex as well as diverse *MKK3* haplotype NILs in RGT Planet background. Error bars are ±SD; a Brown-Forsythe ANOVA test (p < .05, F*(8.000, 4.492) = 33.44, p = .0012) was used to compare the effect of diverse *MKK3* haplotypes on PHS effect. Letters denote significant differences between groups. **F**, PHS effect (G-energy, calculated from PHS data shown in (**D**), see Methods) as a function of α-amylase activity of micro-malted field-grown grains (NZL 2024) with r^2^ value of the linear regression fit in red.

We then explored the impact of different *MKK3* locus haplotypes on GE and GI in a common genetic background. We used marker-assisted breeding to assemble six near-isogenic lines (NIL1–NIL6) in *cv.* RGT Planet by introgressing various *MKK3* locus haplotypes from a range of donors (Supplementary Table S6). We included a FIND-IT premature stop codon variant (MKK3^*270^, i.e., Tryptophan (W) to stop at position 270) in the RGT Planet genetic background (33). In our PHS field trials, a genotype prone to PHS will show a reduction in GE and/or GI in the rain-exposed samples, since re-germination of a pre-sprouted grain is suspended or decelerated, respectively. We analysed germination characteristics of both dry and wet-harvested grain in 2023 and 2024. Assessing the impact of *MKK3* locus haplotype on GE and GI at three weeks post dry harvest (only 2024 season), where dormant cultivars (e.g., *cv.* RGT Planet) still possessed significant dormancy, showed that isolines containing multiple *MKK3* copies or MKK3^Q165^ in single-or multi-copy had no dormancy (Supplementary Fig. S4A, S4B). Critically, the *cv.* RGT Planet knockout variant MKK3^*270^ was fully dormant, supporting *MKK3* as the major dormancy controller (Supplementary Fig. S4A, S4B).

Next, we compared GE and GI in the PHS-inducing wet-harvest grain at 10 weeks (i.e., after dormancy is broken: Fig. 4C; Supplementary Fig. S4C) in each NIL to the dry-harvest samples (2023 and 2024 season). We anticipated a drop in GE and GI after PHS induction. In the wet-harvested samples from 2023, we observed GE’s that were (*i*) unchanged or mildly reduced (i.e., up to 5% in RGT Planet and NIL1 (MKK3^T260^, respectively), (*ii*) strongly reduced (i.e., up to 25% in NIL2 (MKK3^Ref^/MKK3^Ref^) and NIL3 (MKK3^Ref^/MKK3^Ref^/MKK3^Ref^/MKK3^Ref^)) and (*iii*) very strongly reduced (i.e., up to 60% in NIL4 (MKK3^Q165^) and NILs 5 and 6 (MKK3^Ref^/MKK3^Q165^/MKK3^Q165^)) (Fig. 4D,E). We found a similar pattern for GI, with the noticeable difference that multi-copy haplotypes (NIL2/3) are as strongly affected as the NILs with the MKK3^Q165^ variant in single-and multi-copy haplotypes (NIL 4/5/6) (Fig. 4D,E). In the 2024 season, where PHS inducing conditions were suboptimal (Supplementary Fig. S4D-E) we see no significant difference between the dry and PHS harvest germination for the multi-copy haplotypes (NIL2/3; Supplementary Fig. S4F-G). In summary, different GE and GI responses of isolines under both dry and wet harvest conditions confirmed our hypothesis that dormancy is fine-tuned by different *MKK3* haplotypes.

### *MKK3* adaptation to environment, agricultural practice and culture

To understand if haplotype-dependent fine-tuning of dormancy arose through association with environmental, agricultural and/or cultural habits we compared selection pressure between domesticated and wild barleys at the variant level. This showed increase in the frequency of non-synonymous variants in domesticated barley. In contrast, synonymous variant frequencies were not statistically different between wild and domesticated barley (Supplementary Text 1.5). This increased selection of *MKK3* variants in domesticated barley was supported by a median-joining haplotype network analysis of georeferenced wild and domesticated accessions that showed an expansion of haplotype diversity in domesticated barley, whereas wild barley accessions had minimal and dispersed haplotypes (Supplementary Fig. S5; Supplementary Table S8;Supplementary Text 1.5).

We then explored the demographic distribution of selected barley *MKK3* haplotypes at a global scale. Genotypes with single-copy MKK3^T260^ (i.e. unaffected by PHS-inducing conditions) were found in regions exposed to a very high PHS risk due to the co-occurrence of harvest and monsoon weather in the region (Fig. 3A,B; Fig. 4 C-E) (*13*). In contrast, traditional agricultural practices in certain regions of the world (e.g., the highlands of Pamir (Tajikistan, Kyrgyzstan, Pakistan, Afghanistan), the Tibetan plateau (China, Bhutan, Nepal) and Ethiopia favour barleys with no dormancy (*7-9, 34,* 35). Genotypes with the hyperactive Ethiopian MKK3^V79^ variant (Supplementary Text 1.4) and low dormancy match the suitability for double-cropping which increases annual yield and avoids long-term storage and associated pest problems (*7-9, 34*). Tibetan (hull-less) barleys contain many *MKK3* copies and associated lower dormancy in years with high PHS risk (Fig. 2A; Supplementary Fig. S4C; S4F-G; Supplementary Tables S1, S3). In this region, harvest frequently occurs before the grain is fully matured and air-dried prior to processing (roasting and grinding) for human consumption and storage over winter (*35*). Our field experiments showed that multi-copy haplotypes (e.g. NIL2/3) have little grain dormancy, but importantly, that GI was less severely affected by PHS induction (Fig. 4C-E; and Supplementary Figs. S4C, S4F-G). We suggest that a key attribute of the stored seed is the maintenance of vigour that, after planting, is primed to cope with harsh short-season environments allowing most grains to germinate and establish seedlings.

### A hyperactive MKK3 originated in Nordic agriculture

We found multi-copy and hyperactive *MKK3* locus haplotypes in barleys grown in temperate and sub-polar regions (e.g., North America, Scandinavia and the North Atlantic Islands). Re-tracing pedigrees suggested that the MKK3^Ref^/MKK3^Q165^/MKK3^Q165^ haplotype that persists in PHS-susceptible North American malting barleys (Supplementary Table S1)) was derived from the Norwegian *cv.* Domen (*36*) released in the 1950s. Using ddPCR-based genotyping, we traced this Domen haplotype to far older landraces (e.g., Jotun, Asplund, Maskin, Björneby) originating from across Scandinavia, Scotland, the Faroe Islands, Iceland and Russia (Fig. 3A; Supplementary Table S1), where multiple copy *MKK3* locus haplotypes with at least one MKK3^Q165^ variant are found. Extending our haplotype analysis across the *MKK3* locus to a selection of Northern European barley landraces and modern elite varieties using 50K-SNP-array data (*37*) (Supplementary Table S1) revealed that the Scandinavian landraces carrying MKK3^Q165^ cluster together with Bere barleys, the oldest known barley landraces in northern Europe (Supplementary Fig. S6). Bere is still cultivated on the Northern and Western Isles of Scotland where it was introduced more than 5,000 years ago (*38*). ddPCR analyses of 28 different Bere accessions (*39*) revealed the presence of MKK3^Q165^ in all lines alongside variation in copy number (1-4 copies) (Fig. 2A; Supplementary Table S1). Thus, Bere may well be the ancestral Northern European donor of MKK3^Q165^.

In these wet sub-polar regions, barley was selected for quality characteristics that emerge after controlled germination (malting), which is key to the production of alcoholic beverages (beer). Prior quantitative genetic analyses have mapped components of malting quality to the same region as PHS (*18, 26*). Through micro-malting and analysis of *MKK3* NILs 1-6 we found that haplotypes containing at least one hyperactive MKK3^Q165^ (i.e., NILs 4/5/6) have elevated malt quality characteristics including high α-amylase, free limit dextrinase and (1,3;1,4)-β-glucanase enzyme activities (Supplementary Fig. S7A-C). These qualities are in particularly high demand for brewing beer from a mix of barley malt and other starchy grains (*38*). However, these rapid germination characteristics come with the trade-off of increased susceptibility to PHS (Fig. 4F; Supplementary Fig. S7D-E). Interestingly, records from the 18^th^ century describe extinct Norwegian landraces of the ‘Thorebygg’ type as pre-17^th^ century barleys with high sensitivity to moisture during the autumn harvest period (i.e., PHS) (40). These Norwegian landraces were prized for their superior quality characteristics needed for the brewing of traditional farmhouse ales, which used barley malt along with other more widely available starchy cereal grains such as oats, that are normally less suitable for brewing (*40*). If they are as we suspect derived from Scottish Bere barleys it is reasonable to assume that over 1,000 years ago, early Viking farmers would have valued barley with MKK3^Q165^ haplotypes for brewing so highly that they carried grains with them on their travels, developing specific agricultural practices to avoid PHS, such as pre-mature harvest followed by smoke-drying (*42*). They would have been largely responsible for its primary distribution (i.e., within Northern Europe), while a secondary distribution was driven by modern breeding programmes, first successfully to Canada and then to Australia where MKK3^Q165^ failed due to its sensitivity to local climatic conditions, illustrating how a trait with such a strong end use value can be sought and persevere despite a strong risk for crop loss.

## Discussion

The rise of agriculture with its associated domestication of cereal crops initiated arguably the greatest single step development in the history of humans (*1*). Its success required newly domesticated crops to adapt to diverse and changing environmental demands that accompanied primitive cultivation and agricultural expansion. One such demand was the need to match post-harvest dormancy with agricultural practices, environmental conditions and the end use of the crop. By unravelling the complexity of the *MKK3* locus in cultivated barley, we show how it has evolved independently multiple times post domestication to balance dormancy, environmental resilience and critical end-use traits associated with local agricultural practices. Today, more than ever, it is critical that we understand how widespread high-value traits impact a crop’s environmental resilience to avoid high costs from grain loss and quality downgrading.

Here we have assembled evidence that human intervention, enabled by access to environmentally driven variants, shaped the functional landscape of the *MKK3* locus in barley to match end-use requirements. Our data suggest three main driving mechanisms: 1) variation in *MKK3* copy number to adjust transcript abundance, 2) selection and maintenance of *MKK3* amino acid variants with altered kinase activity and 3) distinct combinations of these mechanisms. Genotypes with multiple *MKK3* copies and haplotypes that encode enzymes with higher kinase activity have short dormancy, which increases PHS susceptibility. These have improved end use characteristics that make them suitable either for increasing annual productivity (double-cropping) or improving grain quality characteristics (e.g., short storage and superior malt quality). Barley haplotypes containing MKK3^Q165^, independently selected in Scandinavia and the Caucasus, have generally replaced all other haplotypes in North America for industrial end use while, without current knowledge, providing the lowest climate resilience and highest risk for PHS. Here, we provide a framework to design *MKK3* haplotypes that harmonise regional environments with end use requirements.

A striking observation from our data is the deep historic roots of *MKK3^Q165^*haplotypes. They have been distributed across continents and maintained through millennia despite their associated risk for PHS. This is an example of an extremely valuable trait that persists in compatible regions where grain quality is seldom jeopardised by the environment (high-altitude environments with low humidity such as the plains of Idaho, Montana, Alberta, Saskatoon). Regrettably, its superior and highly desirable malting characteristics also saw *MKK3^Q165^* haplotypes spread to less compatible regions, as evidenced by introgression into AMBA lines grown across North American and into Australian malting barleys (Supplementary Table S1). Such range expansion altered the growing ecosystem to the extent that the trade-off between trait and environment resilience quickly became economically unsustainable.

This study raises an important evolutionary question; is the immense complexity and dynamic nature of the barley *MKK3* locus unique? Between species, MKK3 is a key dormancy regulator across multiple domesticated cereals including rice (14) and wheat (12) that will have responded to similar selection pressures. Within species, the barley pangenome identified more than 150 structurally complex loci (*28*), prompting us to ask how many post-domestication trait-associated loci will exhibit a similarly complex functional haplotype diversity?

A key insight from our study is that the balance between environmental resilience and end use performance of a crop is a delicate one. We show that joint exploration of exotic crop germplasm at the genomic level combined with appropriate phenotypic analyses can provide valuable clues about genetic variants that, when introduced into modern genotypes, may provide a balanced and sustainable solution for the challenges of today.

## Acknowledgments

We would like to extend our gratitude to the following institutions and colleagues for providing the essential seed accessions or DNA sources: the Nordic Genetic Resource Center (NordGen), Sweden; the IPK Genebank at Leibniz Institute of Plant Genetics and Crop Plant Research, Germany; the National Small Grains Collection (NSGC) – GRIN at USDA-ARS, USA; Gary J. Muehlbauer, Kevin P. Smith, Brian J. Steffenson and Shane Heinen from the University of Minnesota, USA; Austin Case from Busch Ag in Ft. Collins, USA; Aaron Beattie from the Crop Development Center at the University of Saskatchewan, Canada; David Moody from Intergrain, Australia; and Blakely Paynter from the Department of Primary Industries and Regional Development, Australia. Special thanks to Hannu Ahokas for discussions on the Finnish barley breeding history. The authors acknowledge Research Computing at the James Hutton Institute for providing computational resources and technical support for the ‘UK’s Crop Diversity Bioinformatics HPC’ (BBSRC grants BB/S019669/1 and BB/X019683/1), use of which has contributed to the results reported within this paper. Maëva Bicard is thanked for her help in extracting weather data.

## Funding

Carlsberg Foundation grant CF15-0236 (B.S).

Carlsberg Foundation grant CF15-0476 (B.S).

Carlsberg Foundation grant CF15-0672 (B.S.).

Carlsberg Foundation grant CF18-0024 (Y.W., Z.H., E.W.).

German Federal Ministry of Education and Research (BMBF) grant 031B0190A (N.S).

German Federal Ministry of Education and Research (BMBF) grant 031B0884A (N.S).

European Union’s Horizon 2020 research and innovation programme grant 862613 (B.J.).

Japan Society for the Promotion of Science (JSPS), KAKENHI, grant JP18H02183 (S.N.).

Biotechnology and Biological Sciences Research Council (BBSRC), grant BB/S004610/1 (M.S., R.W.).

Biotechnology and Biological Sciences Research Council (BBSRC), grant BB/S019669/1 (R.W.). Scottish Government, Rural and Environment Science and Analytical Services Division (RESAS) grant KJHI-B1-2 (M.B., J.R., R.W).

## Author contributions

C.D., M.E.J., and D. V. designed the study. C.D., M.E.J., Y.W., D.V., G.B.F., Z.G., Q.L., S.N., P.R.P., K.S., B.S., N.S., R.W. and E.W. conceived the study. The project and experiments were coordinated by C.D. and M.E.J. The paper was written by M.E.J., C.D., Y.W., R.W., G.B.F., and E.W. *MKK3* genotyping was conducted by Q.L., E.M., J.R., C.V. and C.D. E.M., M.W.R., H.C.T., and C.V. performed the *MKK3* CN analyses (ddPCR). M.E.J., B.J., S.M.K., F.K., Q.L., Y.W., Y.C., F.C., H.D., Z.H., S.T., E.W., M.M. and N.S. carried out the *MKK3* pangenome analyses. *MKK3* pantranscriptome analyses were performed by M.E.J., M.B., Q.L., M.S., and R.W. MKK3 *in vitro* kinase assays and protein modelling were managed by M.E.J. and J.A.C.S. Breeding and field work for *MKK3* NILs were supervised by D.V., A.B., P.M.L.R., A.L., M.T.S.N., P.A.P., N.W.T. and C.D. The *MKK3* FIND-IT analysis was conducted by M.W.R., and C.V., M.E.J., K.B.B., E.M. and C.V. oversaw the *MKK3* germination assays, while K.B.B. performed malt analyses. All authors read and provided valuable input on the manuscript.

## Competing interests

M.E.J., C.B.A., K.B.B., J.A.C.S, Z. G., S.M.K., F.K., Q.L., M.T.S.N., P.R.P., M.W.R., H.C.T., E. M., N.W.T., S.T., C.V., B.S. and C.D. are current or former employees of the Carlsberg Research Laboratory. All other authors declare no competing interests.

## Data and materials availability

All data are available in the main text or the supplementary materials.

## Supplementary Materials

Materials and Methods

Supplementary text Figs. S1 to S7

Tables S1 to S8

References*43*–*62* are only cited in the materials and methods

