## Supplementary methods and materials for "Post-Domestication selection of MKK3 Shaped Seed Dormancy and End-Use Traits in Barley"

[1]Carlsberg Research Laboratory, J.C. Jacobsens Gade 4, 1799 Copenhagen V, Denmark. [2]SECOBRA Recherches; Centre de Bois Henry, 78580 Maule, France.[3]Group of Alpine Paleoecology and Human Adaptation (ALPHA), State Key Laboratory of Tibetan Plateau Earth System, Environment and Resources (TPESER), Institute of Tibetan Plateau Research, Chinese Academy of Sciences; Beijing 100101, China. [4]Department of Genetics, University of Cambridge, Cambridge; CB2 3EH, UK. [5]Ancient Environmental Genomics Initiative for Sustainability, Globe Institute, University of Copenhagen, 1350, Copenhagen, Denmark. [6]Centre for Ancient Environmental Genomics, Globe Institute, University of Copenhagen; Copenhagen, Denmark. [7]International Barley Hub (IBH)/James Hutton Institute (JHI); Errol Road Invergowrie Dundee DD2 5DA Scotland. [8]University of Chinese Academy of Sciences; Beijing 100049, China. [9]Key Laboratory of Western China's Environmental Systems (Ministry of Education), College of Earth and Environmental Sciences, Lanzhou University; Lanzhou 730000, China. [10]School of Agriculture, Food and Wine, University of Adelaide, Waite Campus; Urrbrae, SA 5064, Australia. [11]University Zürich; Zollikerstrasse 107, 8008 Zürich, Switzerland. [12]Leibniz Institute of Plant Genetics and Crop Plant Research (IPK) Leibniz Institute of Plant Genetics and Crop Plant Research (IPK); 06466 Stadt Seeland, Germany. [13]Division of Crop Design Research, Institute of Crop Science, NARO; Tsukuba, 305-8518, Japan. [14]Okayama University; Kurashiki, 710-0046, Japan. [15]Setsunan University, Faculty of Agriculture; Hirakata, 573-0101, Japan. [16]Kazusa DNA Research Institute, Department of Frontier Research and Development; Kisarazu, 292-0818, Japan. [17]Crop Plant Genetics, Institute of Agricultural and Nutritional Sciences, Martin Luther University Halle-Wittenberg; Halle (Saale), Germany. [18]Division of Plant Sciences, School of Life Sciences, University of Dundee; Dundee, UK, †Current address, Novo Nordisk A/S; 2880 Bagsværd, Denmark. ‡Current address, Crop Genetics and Biotechnology Department of Agroecology, Aarhus University; Slagelse, Denmark. §Current address, DLF Seeds A/S; 4000 Roskilde, Denmark. ¶These authors contributed equally: Morten E. Jørgensen, Dominique Vequaud, Yucheng Wang. *Corresponding authors..

**The PDF file includes:**

Materials and Methods
Supplementary Text
Figs. S1 to S7
Tables S1 to S8
References

### Materials and Methods

*MKK3* barley pangenome (BPGv2) assembly analyses

The *MKK3* gene copy representative (HORVU.MOREX.PROJ.5HG00474740) was used to Blast against all 76 pangenome assemblies in the extended barley pangenome (BPGv2) (*28*). Full-length sequences with identity over 95% were extracted and used for further analyses. All sequences were aligned using MAFFT v7.490 (*43*). Variations among *MKK3* gene copies at genomic DNA, CDS and respective protein level were collected, and *MKK3* haplotypes in each pangenome accession were summarised using R v4.2.2 (*44*).

*MKK3* barley pan-transcriptome (BPTv1) analyses

*MKK3* transcript abundance was extracted from the pan-transcriptome data (BPTv1) (*29*). Briefly, annotation was identified both for the genotype-specific reference transcript datasets (Supplemental Table S5) and the pan-transcriptome reference transcript dataset (PanBaRT20: chr5H56507 and chr5H56511). The pan-transcriptome comprises 19 domesticated and 1 wild barley accessions, which are a subset of BPGv2 (*28*). RNA-seq for five different tissues (Shoot, Root, Caryopsis, Inflorescence, Embryonic) and three replicates for each accession was generated. For gene expression, the RNA-seq was mapped against the PanBaRT20 reference transcript dataset, and transcript per million (TPM) values were extracted.

*MKK3* barley BARN analyses

Gene expression for the 209 European two-rowed spring malting barleys was extracted from the published BARN dataset (*45, 46*) using the gene identifier BaRT2v18chr5HG281230 for *MKK3* and plotted against the *MKK3* CN (Supplementary Table S7).

*MKK3* cluster structure in BPGv2

Based on the reference genome annotation, 15 gene sequences on each side of the *MKK3* gene copy (HORVU.MOREX.PROJ.5HG00474740) were extracted from all 76 pangenome assemblies (*28*). The 31 genes were blasted against the 76 *de novo*-assembled genomes using NCBI-Blast (word_size of 11 and perc. identity of 90, other parameters as default) (*47*). Plots were generated from the Blast result coordinates by scaling based on the mid-point between HORVU.MOREX.r3.6HG0617300/HORVU.MOREX.PROJ.6HG00545250 and HORVU.MOREX.r3.6HG0617710/ HORVU.MOREX.PROJ.6HG00545670. All Blast results in the region (+/- 1Mb) around this mid-point were plotted using R-base (44).

*MKK3* promoter structure in BPGv2

2 Kbp upstream of the ATG start site of all *MKK3* copies across the 76 pangenome assemblies (28) were aligned and pairwise comparisons performed using CLC Main Workbench (v22.02, Qiagen).

Synteny analysis of *MKK3* chromosomal region in BPGv2

The *MKK3* region from the reference genome assembly of *cv.* Morex (48) (MorexV3) was identified by searching the *MKK3* genes against the assembly using Blastn. For synteny analysis, chromosome 5H (chr5H) of BPGv2 assemblies was extracted and aligned against the chr5H from the MorexV3 using minimap2 (V2.24) (*49*). The outputs of minimap2 were imported in R statistical environment, and alignments of more than 2Kb sizes from a region 583.516–587.595 Mb were extracted and used for creation of x-y plot.

*MKK3* haplotype network analysis in BPGv2

*MKK3* gene sequences (ATG to STOP including introns) identified above were aligned using CLC Main Workbench (v22.02, Qiagen). Two sequences were removed for this analysis due to major deletions in their sequences that affected the analysis: 1) contig_corrected_v1_3602_HORVU.HOR_8148.CNS.contig_corrected_v1_3602G00694700; and 2) contig_corrected_v1_3602_HORVU.HOR_8148.CNS.contig_corrected_v1_3602G00694750). Median-joining haplotype network was generated using PopART (50) with an epsilon value of 0.

*MKK3* CN estimation in BPGv2

CN estimation was carried out in the BPGv2 (28) associated dataset (PRJEB53924) that comprises 1,000 plant genetic resources and 315 elite cultivars as well as the BARN dataset (45, 46) and the exome capture dataset (PRJEB8044) that represents a globally distributed diversity panel of barley accessions (56). *K*-mers (k=21) were first generated from the $MKK3^{Ref}$ gene (HORVU.MOREX.PROJ.5HG00474740) using jellyfish v2.2.10 (*51*). The *k*-mer list was then cleaned up using the MorexV3 HiFi assembly (48) to remove *k*-mers from repetitive regions. All *k*-mer counts were normalised to counts from the *cv.* Morex sequence from the short read based pangenome (BPGv1) (*52*). For each accession, *MKK3* CN was estimated as the median count of all *k*-mers from this accession. All *k*-mer counting was done using Seal (*53*) (BBtools). Data normalisation and summary was carried out in R v4.2.2 (*44*). CN variation was illustrated using QGIS Geographic Information System (http://www.qgis.org).

Targeted *MKK3* read enrichment and genotyping in barley

To investigate polymorphism at the *MKK3* locus, we implemented a targeted read enrichment and genotyping pipeline optimized for regions with duplicated genomic sequences. This approach enhances sensitivity for low-frequency variants that are frequently overlooked in conventional genome-wide analyses. We analyzed two publicly available datasets. The exome capture dataset (PRJEB8044) represents a globally distributed diversity panel of barley accessions (45, 46). The BPGv2 associated dataset (PRJEB53924) comprises 1,000 plant genetic resources and 315 elite cultivars. Sequencing reads were quality-trimmed using Cutadapt (54) (version 4.2, Python 3.9) to remove adapter sequences and low-quality bases, retaining reads of 30 bp or longer. For targeted enrichment, 31-mers were generated from the full-length *MKK3* gene (7,434 bp) from HORVU.MOREX.r3.5HG0537440 using KMC tools (55) (v3.2.1). To minimize cross-mapping between paralogs, the homologous copy HORVU.MOREX.r3.5HG0537390 was masked in the MorexV3 chromosome 5H reference (48). We then applied kmc_tools simple intersect -cx1 to subtract k-mers found in the masked genome, retaining only those unique to *MKK3*. These unique k-mers were exported using kmc_dump and converted to fasta format. All paired-end reads were scanned for matches to *MKK3*-unique k-mers. Read pairs containing at least one match were retained. The sample-enriched reads were aligned to the masked chromosome 5H reference using BWA (56) (v0.7.17).

Alignment files were sorted and deduplicated with SAMtools (56) (v1.15). To identify all possible coding and intronic variants, we generated a synthetic VCF representing every potential single-nucleotide substitution in the *MKK3* coding region, yielding 22,302 variant positions. Genotyping was performed using bcftools (56) mpileup and call (v1.15), and genotype (GT) and read depth (DP) fields were parsed using custom Python and R scripts. Global *MKK3* haplotype variation was illustrated using QGIS Geographic Information System (http://www.qgis.org). Annual mean precipitation (WorldClim2.1 variable BIO12 (30)) is shown as red gradient (spatial resolution 5 min).

MKK3 *in vitro* kinase assays

All assays were carried out by Reaction Biology (Reaction Biology Europe GmbH). cDNA sequences of $MKK3^{Ref}$ (reference (HORVU.MOREX.PROJ.5HG00474740) and point mutants $MKK3^{V79}$, $MKK3^{Q165}$, $MKK3^{T260}$ and $MKK3^{N383}$ as well as the kinase dead substrate $AtMPK1^{K61R}$ were cloned into an IPTG (Isopropyl-β-D-thiogalactopyranoside)-inducible plasmid for recombinant bacterial expression. The *MKK3* open reading frame was cloned in-frame to an N-terminal GST-tag sequence (GST, glutathione-S-transferase, S. japonicum) to enable purification by GST-GSH affinity chromatography. Chemically competent *E.coli* BL21 (strain Codon+) were transformed with the respective IPTG-inducible *MKK3* expression plasmids and plated under antibiotic (ampicillin) selection. Single cultures were inoculated into 27 °C over-night cultures supplemented with 10 μM of IPTG (Carl Roth, Karlsruhe, Germany, #CN08.3) to induce protein expression.

Bacteria were lysed under native conditions by sonication in HEPES (Carl Roth, Karlsruhe, Germany, #9105.4) pH 7.5, 150 mM NaCl (Carl Roth, Karlsruhe, Germany, #P029.2), 1 mM EGTA (Carl Roth, Karlsruhe, Germany, #3054.3), 2 mM EDTA (Carl Roth, Karlsruhe, Germany, #8043.2) , 2 mM DTT (Carl Roth, Karlsruhe, Germany, #6908.2), protease inhibitor cocktail (SERVA Electrophoresis GmbH, Heidelberg, Germany #39106.53), 1 mg/ml lysozyme (Carl Roth, Karlsruhe, Germany #8259.2), 5 μg/ml DNAse A (Sigma-Aldrich/Merck, Darmstadt, Germany #DN25). Cellular debris and the insoluble fraction were removed by centrifugation. The cleared lysate was applied to a GSH-affinity (Glutathione) matrix (Sigma-Aldrich/Merck, Darmstadt, Germany, #64510), GST-tagged protein was bound, and the beads were washed 3 times prior to elution by free reduced glutathione (Carl Roth, Karlsruhe, Germany #6382.2). The concentrations of the eluted proteins were determined using the Bradford method (reference protein: bovine serum albumin (Sigma-Aldrich/Merck, Darmstadt, Germany, #A-7638). The final buffer composition of the protein preparation is 50 mM HEPES pH 7.5, 100 mM NaCl, 5 mM DTT, 15 mM reduced glutathione, 20% (v/v) glycerol (Carl Roth, Karlsruhe, Germany, #3783.2). Proteins were aliquoted, snap-frozen and stored at -80 °C.

Recombinant MKK3 was incubated with the upstream activating kinase MLK4 (MLK4, Lot002 (Reaction Biology Europe, #1078-0000-1)) at a concentration of 200 μg/ml for MKK3 and 20 μg/ml for MLK4 in kinase assay buffer 70 mM HEPES pH 7.5, 5 mM $MgCl_2$ (Sigma-Aldrich/Merck, Darmstadt, Germany, #M0250), 3 μM Na-orthovanadate (Sigma-Aldrich/Merck, Darmstadt, Germany, #S6508), 1.2 mM DTT, 50 μg/ml PEG20000 (SERVA Electrophoresis GmbH, Heidelberg, Germany, #33138), 100 μM ATP (Promega, Walldorf, Germany, #V9101) for 60 minutes at 30 °C. The activation mix was diluted into the $MPK1^{K61R}$ phosphorylation reaction to final concentrations of 2 μg/ml MLK4, 20 μg/ml MKK3 and 200 μg/ml MPK1 K61R. The ATP concentration was adjusted to 5 μM and radioactive $^{33}$P-γ-ATP (5E+05 cpm/μl)

(Hartmann Analytic, Braunschweig, Germany, #FF301T) was added to enable visualisation of AtMPK1$^{K61R}$ phosphorylation. The total assay volume was 50 µl. The final assay contained 70 mM HEPES-NaOH pH 7.5, 5 mM MgCl2, 3 µM Na-orthovanadate, 1.2 mM DTT, 10 µM ATP/33P-γ-ATP (approx. 2.5 x $10^7$ cpm per sample), kinase and substrate. The reaction cocktails were incubated at 30 °C for 60 minutes. The reaction was stopped with 20 µl LDS-Gel loading buffer (Invitrogen/ThermoFisher Scientific, Freiburg, Germany, #NP0007) and incubated for 3 minutes at 95 °C. 20 µl of the mixture were loaded on a 4-12% Bis-Tris-SDS-poly acrylamide gel (Invitrogen/ThermoFisher Scientific, Freiburg, Germany, #NP0321). The samples were analysed by electrophoresis. After removal of the radioactive ATP band, the gel was Coomassie-stained, dried and exposed to X-ray film (Carestream Health Deutschland, Stuttgart, Germany, #8736936). Absolute phosphorylation was quantified by densitometric analysis using GelAnalyser2010 software (Gelanalyser.com; raw volume with background subtracted).

NILs breeding

Near-isogenic lines (NILs) harbouring different *MKK3* haplotypes identified by ddPCR genotyping or in exome sequencing of genomic resources (*56*) were derived from crosses between donor genotypes (Supplementary Table S6) and the recurrent parent RGT Planet. After two cycles of backcross and one selfing step ($BC_2S_1$), homozygous plants at the *MKK3* locus were retrieved. 8 NILs were developed (Supplementary Table S6). Plants were grown in a greenhouse at 18°C under 16/8-hour light/dark cycles. Foreground and background molecular markers were used in each generation to assist plant selection. Respective $BC_2S_1$ plants were genotyped and grown to maturity. Grains were harvested and further propagated in field plots and harvested and threshed using a ZURN 150 plot combine (ZURN, Germany), and sorted by size (threshold, 2.5 mm) using a 'Blue Rocket' Pfeuffer SLN3 sample cleaner (Pfeuffer GmbH, Germany).

FIND-IT variant identification

Barley FIND-IT variant MKK3$^{*270}$ [ID# CB-FINDit-Hv-039] was identified and isolated in the RGT Planet variant library as described (33) and backcrossed with the recurrent parent RGT Planet as described for NILs.

Field experiments and PHS-testing setup

*Cv.* RGT Planet, NILs and the FIND-IT variant MKK3$^{*270}$ were planted in a field mini-plot (1 $m^2$) in France. The first set (from here on referred to as 'dry') was harvested at maturation and only exposed to rain. After harvest of the first set, the second set (referred to as 'wet') was harvested after 30 days of natural irrigation (2023 season) or a combination of man-made and natural irrigation (2024 season). All grains were dried and analysed after ripening had naturally broken RGT Planet dormancy (10 weeks after harvesting).

Grain germination

100 grains of each barley line were germinated in duplicates in Petri dishes on 2 layers of Whatman paper (#1) and 4 ml of distilled water for 72 hours at 20 °C. After 24, 48 and 72 hours, samples were visually assessed for percentage of germinating grains using manual inspection (2020-2023 data) or via automated imaging and analysis (Reshape Biotech, Denmark). The germination energy (GE) describes the percentage of germinated kernels and is calculated by

summing germinated kernels every 24 hours for 3 days. The germination index (GI) is an indicator of the germination speed throughout a period of 3 days, described with the following equation: 10∗(24h+48h+72h)/(24h+2∗48h+3∗72h).

Micro-malting and hydrolytic enzyme activity

Non-dormant barley samples of *MKK3* haplotype NILs 1-6 and *cv.* RGT Planet were micro-malted in perforated stainless-steel boxes. For steeping, barley samples were placed in individual containers (three repetitions with 100 g each, graded >2.5 mm) and submerged in 16 °C fresh water to reach 33% moisture content on day 1, and 43% moisture content on day 2. The actual water uptake of individual samples was determined as the weight difference between initial water content, measured with Foss 1241 NIT instrument (Foss A/S, Denmark), and the sample weight after surface water removal. Following the last steep, the barley samples were maintained at a steeping degree of 43% and were germinated for 4 days at 16 °C. After each 24 hours, samples were checked for moisture content and sprayed with additional water to overcome possible respiration loss. After the germination process, the barley samples were kiln-dried in a Curio kiln using a two-step ramping profile. First ramping step started at a set point of 27 °C and a linear ramping at 2 °C/h to the breakpoint at 55 °C using 100% fresh air. Second linear ramping was at 4 °C/h, reaching a maximum of 85 °C. This temperature was kept constant for 90 minutes using 50% fresh air. The kiln samples were cured using a manual root-removal system from Wissenschaftliche Station für Brauerei, Munich, Germany. α-amylase activity was measured using the Ceralpha method (Ceralpha Method MR-CAAR4, Megazyme) modified for Gallery Plus Beermaster (Thermo Fisher Scientific). β-glucanase activity was measured using the Malt β-Glucanase/Lichenase Assay Kit (MBG4 Method) modified for Gallery Plus Beermaster (Thermo Fisher Scientific). Free limit-dextrinase activity was measured using the Pullulanase/Limit-Dextrinase Assay Kit (PullG6 Method).

Weather data

Weather data was downloaded from the Agri4Cast resources portal Agri4Cast Data (europa.eu) for the years 2023 and 2024.

50K-SNP-array cluster analysis

Marker data from the *MKK3* region (5H:580-588Mb, upstream 5M to the end of the chromosome) was extracted from published 50K-SNP-array data (37) of 391 samples and cleaned for further analyses. Pair-wise genetic distance between individuals was calculated as the average of their per-locus distances in this region using R package stringdist. Hierarchical clustering was done using R base function ‘hclust’ based on this genetic distance matrix.

Droplet digital PCR (ddPCR) copy number (CN) variation analysis

CN variations were assessed in DNA samples using ddPCR according to Bio-Rad instructions. The ddPCR assay was performed using a reaction mixture composed of 4× ddPCR Multiplex Supermix (Bio-Rad, Hercules, CA, USA), with primers and probes added at concentrations of 900 nM and 250 nM, respectively. A volume of 5 μl of diluted DNA was included in the reaction mixture, and the remaining volume was made up with molecular-grade water to achieve a total reaction volume of 20 μl. The preparation of the reaction mixture was conducted on a QX200

AutoDG Droplet Digital PCR system (Bio-Rad, Hercules, CA, USA). In this process, each 20 µl of reaction mixture was partitioned into droplets using an oil film. Subsequently, PCR amplification was performed using an Uno96 thermal cycler (VWR, Radnor, PA, USA) with the following temperature profile: initial enzyme activation at 95 °C for 10 minutes, followed by 40 cycles of denaturation at 94 °C for 30 seconds, annealing/extension at 55 °C for 1 minute, and final enzyme deactivation at 98 °C for 10 minutes. All temperature transitions were executed at a rate of 2 °C/s. The ddPCR reactions were analysed using a QX600 Droplet Reader (Bio-Rad, Hercules, CA, USA), and the resulting data were subsequently processed using Bio-Rad QX Manager software (Bio-Rad, Hercules, CA, USA). For *MKK3* copy number analysis, we employed a tailored ddPCR assay to differentiate between copies of $MKK3^{E165}$ and $MKK3^{Q165}$ (Unique BioRad assay ID: dMDS533374121). For reference purposes, we employed a reference assay with the following sequences: Forward primer 5'-AAT GGT ACG AGA GGA AGG-3', Reverse primer 5'-GGC ATA GCT TGT CCC A-3' and Probe 5'-Cy5.5-ATC ACT GGT CAG AGG GT-3'Iowa Black RQ-Sp.

MKK3 protein structure modelling and visualisation

A structural model of the reference MKK3 from *cv.* Morex (28) (HORVU.MOREX.PROJ.5HG00474740) was retrieved from the Alphafold2 database at EMBL-EBI (https://alphafold.ebi.ac.uk/entry/A0A140JZ20) which was also found in the Alphafill database (https://alphafill.eu/model?id=AF-A0A140JZ09-F1-model_v4&identity=30). At 30% identity, it was possible to transplant ATP (from the structure with PDB-ID: 7F2X) into the ATP binding pocket of MKK3. The protein and amino acids of interest were visualised using PyMol 2.5.5 (Schrödinger LLC, USA).

### Supplementary Text

Supplementary text 1.1. Haplotype specific *MKK3* expression levels

We explored differences in *MKK3* transcript abundance in genotypes from the barley pan-transcriptome BPTv1 (*29*). Barley landraces with single copy MKK3$^{REF}$ haplotype (e.g. HOR13942) have moderate *MKK3* expression across all tissues, in the same range as wild barley with the MKK3$^{S76+N383+M394}$ haplotype B1K-04-12 [FT11] and East Asian barleys with MKK3$^{T260}$ haplotype (e.g. Akashinriki) (Fig. 2B), known to have high grain dormancy (*13*). European modern cultivars and landraces with single copy MKK3$^{R350+N383}$ (e.g. RGT Planet and HOR 13821) have increased *MKK3* expression (Fig. 2B), correlating with a reduction in grain dormancy (*13*). Interestingly, we observe that aligned promoter sequences from MKK3$^{R350+N383}$, MKK3$^{T260}$ or MKK3$^{S76+V79+N383+M394}$ haplotypes are largely identical within their haplotype groups, whereas MKK3$^{REF}$ haplotypes with similar expression levels show low conservation (Supplementary text Figure 1.1A). Inversions did not influence expression in single-copy haplotypes (MKK3$^{T260}$; MKK3$^{R350+N383}$; Fig. 2B; Supplementary Fig. S2).

In *MKK3* multi-copy lines increased transcript levels were observed. Morex with two *MKK3* gene copies (MKK3$^{Ref}$/MKK3$^{Ref}$) and HOR7552 with four *MKK3* gene copies (MKK3$^{Ref}$/MKK3$^{Ref}$/MKK3$^{Ref}$/MKK3$^{Ref}$) have approximately two to four times higher *MKK3* expression than accessions with a single MKK3$^{REF}$-copy, respectively. Similarly, when comparing expression and *MKK3* copy number across transcriptomes of 209 European two-rowed spring malting barleys in BARN (45, 46), we observe increased *MKK3* expression with increasing copy number (Supplementary text Figure 1.1B).

Two accessions, modern US *cv*. Hockett (MKK3$^{REF}$/MKK3$^{Q165}$) and landrace OUN333 (MKK3$^{R350+N383}$/MKK3$^{REF}$) from Nepal, also carry two copies of *MKK3*, however both lines have higher expression than *cv*. Morex (Fig. 2B). Importantly, ddPCR based copy number analysis of *cv*. Hockett indicates three *MKK3* copies (MKK3$^{Ref}$/MKK3$^{Q165}$/MKK3$^{Q165}$) for the seed stock tested, whereas the BPGv2 genome assembly, derived from a different seed stock, shows two copies (MKK3$^{Ref}$/MKK3$^{Q165}$). This result is in agreement with our CN analyses of seed stocks of Scandinavian landraces and North American AMBA lines where we also find diverse multi-copy MKK3$^{Q165}$ haplotypes for the same accession from different genebanks or even within the same accession batch (Fig. 1B, Supplementary Table S1).

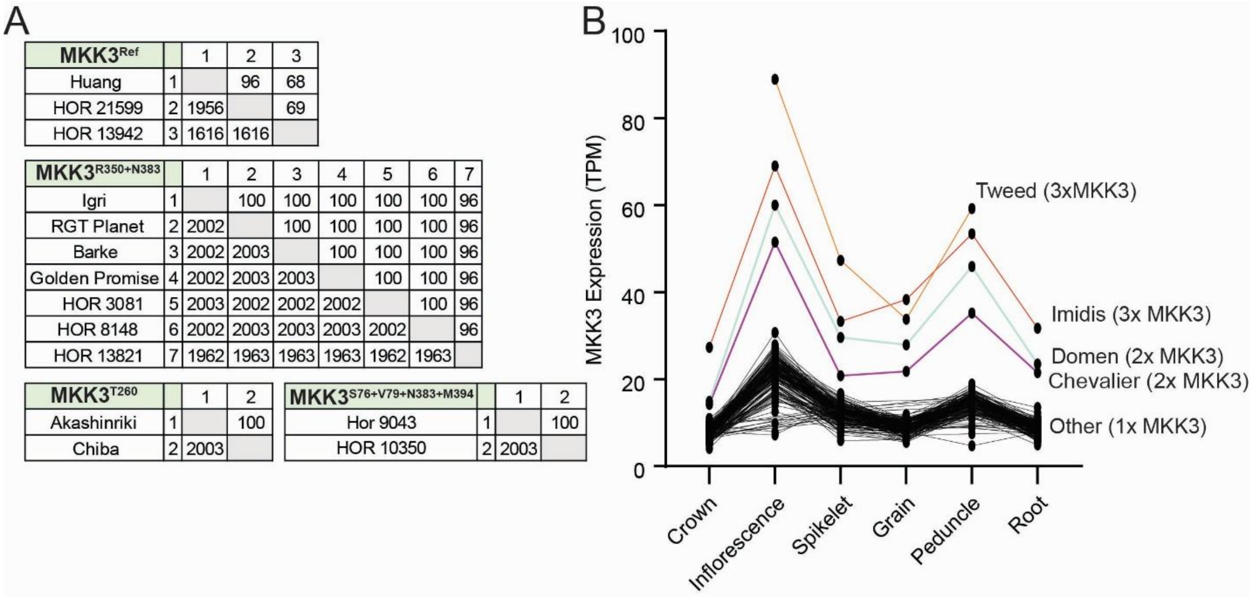

**Supplementary text Figure 1.1. Promoter analysis and MKK3 expression correlation to CNV. A**, Pairwise comparison of the 2 kb promoter segment of *MKK3* in assemblies carrying *MKK3* haplotypes $MKK3^{Ref}$, $MKK3^{R350+N383}$, $MKK3^{T260}$ and $MKK3^{S76+V79+N383+M394}$ (upper and lower comparison is % identity and identities of 2003 bp, respectively). **B**, *MKK3 k*-mer-based copy number estimation and expression data across six tissues and 192 European two-rowed spring barley in BARN (*46*). Accessions with >1 *MKK3* copy number are highlighted with coloured lines, MKK3 copy number per accession is indicated in brackets.

### Supplementary text 1.2. *MKK3* in *Hordeum bulbosum*, a crop wild relative to barley

The wild species *Hordeum bulbosum* is the closest perennial relative of barley that separated from barley some 4,5 million years ago. Thus, *Hordeum bulbosum* is considered the secondary gene pool of cultivated barley (57). Here, we investigated MKK3 amino acid haplotypes in the recently resolved pangenome of multiple, diverse *Hordeum bulbosum* clones (31) relative to the barley genomic reference HORVU.MOREX.PROJ.5HG00474740 ($MKK3^{Ref}$). We find four amino acid changes (i.e. $MKK3^{L73}$, $MKK3^{R177}$, $MKK3^{S313}$ and $MKK3^{N383}$) that exist across all *H. bulbosum* clones, suggesting that variation at these positions occurred after the split of *Hordeum bulbosum* and barley (Supplementary text Table 1.2.1). Interestingly, $MKK3^{N383}$ and $MKK3^{L73}$ variants are also found with the domesticated barley gene diversity space suggesting that variation at these positions are under continues selection and that they arose recently in the barley reference genome cultivar *cv.* Morex (cv. Morex has the alternate alleles $MKK3^{P73}$ and $MKK3^{D383}$).

**Supplementary text Table 1.2.1.** MKK3 copy number and amino acid haplotype variation in the *Hordeum bulbosum* pangenome .

| Name | Ploidy | Gene | AA Hap variation |
|---|---|---|---|
| CP1_HORVU.MOREX.r3.5HG0537390.ORF | | REF | REF |
| H. bulbosum A17 | Tetraploid | HBULBOSUM.A17.PROJ.r1.5H_1G00847750.1 | L73+I109+R177+S313+N383 |
| | | HBULBOSUM.A17.PROJ.r1.5H_2G00905190.1 | L73+I109+R177+S313+N383 |

| | | | |
|---|---|---|---|
| | | HBULBOSUM.A17.PROJ.r1.5H_3G00965960.1 | L73+I109+R177+S313+N383+P523 |
| | | HBULBOSUM.A17.PROJ.r1.5H_4G01023630.1 | L73+I109+R177+S313+N383+P523 |
| H. bulbosum A40 | Tetraploid | HBULBOSUM.A40.PROJ.r1.5H_1G00867260.1 | L73+I109+R177+S313+N383+S514 |
| | | HBULBOSUM.A40.PROJ.r1.5H_1G00867520.1 | L73+R177+D224+S313+N383 |
| | | HBULBOSUM.A40.PROJ.r1.5H_2G00923560.1 | L73+R177+D224+S313+N383+S313+N383 |
| | | HBULBOSUM.A40.PROJ.r1.5H_4G01035670.1 | L73+R177+S313+A341+N383 |
| H. bulbosum A42 | Tetraploid | HBULBOSUM.A42.PROJ.r1.5H_1G00856290.1 | L73+R177+S313+ N383+A394 |
| | | HBULBOSUM.A42.PROJ.r1.5H_2G00912640.1 | L73+R177+S313+ N383+G514 |
| | | HBULBOSUM.A42.PROJ.r1.5H_3G00965830.1 | L73+R177+S313+ N383+D224 |
| | | HBULBOSUM.A42.PROJ.r1.5H_4G01021090.1 | L73+R177+S313+ N383+S514+del523 |
| H bulbosum FB19_001 | Tetraploid | HBULBOSUM.FB19_001_1.PROJ.r1.5H_1G00925660.1 | L73+R177+S313+N383 |
| | | HBULBOSUM.FB19_001_1.PROJ.r1.5H_2G00984150.1 | L73+R177+S313+N383 |
| | | HBULBOSUM.FB19_001_1.PROJ.r1.5H_3G01043600.1 | L73+R177+S313+M345+N383+D433+del523 |
| | | HBULBOSUM.FB19_001_1.PROJ.r1.5H_4G01102400.1 | L73+R177+S313+N383 |
| H. bulbosum FB19_011_3 | Diploid | HBULBOSUM.r1.5H_1G00923610.1 | L73+R117+S313+N383+S388 |
| | | HBULBOSUM.r1.5H_2G01033500.1 | L73+R117+S313+N383 |
| H. bulbosum FB19_028_3 | Tetraploid | HBULBOSUM.FB19_028_3.PROJ.r1.5H_1G00884650.1 | L73+R177+S313+N383+N242 |
| | | HBULBOSUM.FB19_028_3.PROJ.r1.5H_2G00944860.1 | L73+R177+S313+N383 |
| | | HBULBOSUM.FB19_028_3.PROJ.r1.5H_3G01003130.1 | L73+R177+S313+M345+N383+D433+del523+N242 |
| | | HBULBOSUM.FB19_028_3.PROJ.r1.5H_4G01060400.1 | L73+R177+S313+N383 |
| H. bulbosum FB20_005_1 | Diploid | HBULBOSUM.FB20_005_1.PROJ.r1.5H_1G00499480.1 | L73++V105+R177+S313+N383 |
| | | HBULBOSUM.FB20_005_1.PROJ.r1.5H_2G00559140.1 | L73++V105+R177+S313+N383+S388 |
| H. bulbosum FB20_029_7 | Diploid | HBULBOSUM.FB20_029_7.PROJ.r1.5H_2G00553450.1 | L73+R177+S313+N383 |
| | | HBULBOSUM.FB20_029_7.PROJ.r1.5H_2G00553730.1 | L73+R177+S313+N383+S388 |
| H. bulbosum GRA2256_1 | Tetraploid | HBULBOSUM.GRA2256_1.PROJ.r1.5H_2G00876990.1 | L73+H149+R177+L217+S313+N383+A394+L495 |
| | | HBULBOSUM.GRA2256_1.PROJ.r1.5H_4G00984250.1 | L73+I109+R177+S313+N383+A493+del523 |
| H. bulbosum PI365428 | Diploid | HBULBOSUM.PI365428.PROJ.r1.5H_1G00477900.1 | L73+R177+S313+N383 |
| | | HBULBOSUM.PI365428.PROJ.r1.5H_2G00535950.1 | L73+R177+S313+N383 |

We next investigated the occurrence of CNV in the *Hordeum bulbosum* pangenome clones. Here, three clones (A40, GRA2256-1 & FB20-029-7) show translocation and/or deletions of *MKK3* across sub-genomes (Supplementary text Fig. 1.2.1*)*.

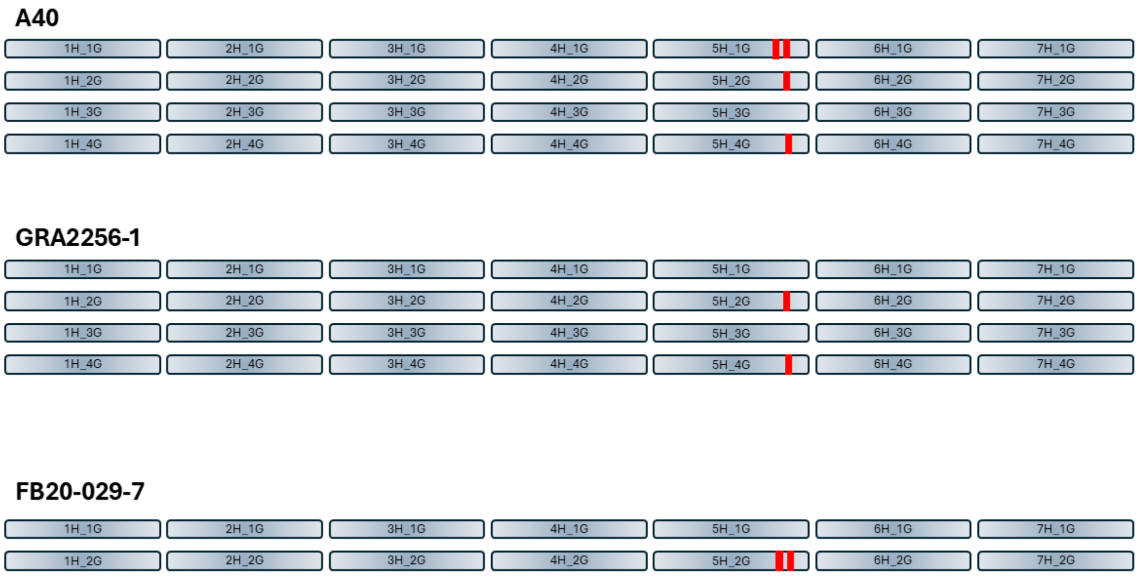

**Supplementary text *Fig. 1.2.1.*** Schematic overview of three *Hordeum bulbosum* clones with uneven distribution of *MKK3* genes among sub-genomes, exemplifying the heterozygosity that comes with its outcrossing nature. Grey bars represent chromosomes, red lines represent the *MKK3* gene. Shown clones are A40 (Tetraploid; 2n4x=28) with two copies on the 5H_1G chromosome, one copy on 5H_2G, none on 5H_3G and one on 5H_4G; GRA2256-1 (Tetraploid; 2n4x=28) lacking *MKK3* on 5H_1G and 5H_3G, while having one copy on 5H_2G and 5H_4G; and FB20-029-0 (Diploid; 2n2x=14) with two copies on 5H_2G and none on 5H_1G.

### Supplementary text 1.3. The barley MKK3$^{Q165}$ variant in a structural context

A MKK3 protein model indicates that E165 is hydrogen bonded, via the main chain, to the adenine of the ATP substrate, controlling its position and orientation and thus, indirectly, the position of the gamma phosphate of ATP. Therefore, E165 possibly affects the orientation of the adenine ring and of the hinge loop between the N-lobe and the C-lobe of the kinase. In addition, the E165 sidechain is salt bridged to K226, E147, and part of a larger salt bridge network surrounding K226. Removing the negative charge in E165 by mutation to Q may increase the flexibility of the 165-167 loop and allow displacement of the ATP (away from K226). At the same time, releasing this salt bridge could influence the position and flexibility of the so-called DFG loop nearby (D229, F230 and G231) (*32*). The highly conserved DFG loop is in contact with the phosphates of ATP and is known to alternate between "active" or „Asp-in" and "inactive" or „Asp-out" conformations (*32*). A change of the evolutionary conserved E165 to Q165 could favor the active „Asp-in" conformation of MKK3, resulting in increased kinase activity.

### Supplementary text 1.4. An Ethiopian *MKK3* haplotype associated with low dormancy shows increased MKK3 kinase activity *in vitro*

In the highlands of Ethiopia barley is grown predominantly for food but also for malting and traditional beverage production. Here, low dormancy levels important for double cropping have been reported (*34*). We identified a MKK3$^{V79}$ variant in 43/83 Ethiopian landraces from a collection of 1,707 diverse barley lines (Supplementary Table S3). *In vitro* kinase activity measurements showed that MKK3$^{V79}$ is a kinase with increased activity compared to MKK3$^{Ref}$ (Supplementary Fig. S3D). This suggests that this unique variant found in Ethiopian *MKK3* haplotypes is causative for the low dormancy trait important for environmental adaptation and agricultural traditions. In PHS trials (FRA 2023) we compared GE and GI in the PHS-inducing wet-harvest grain at 10 weeks (i.e., after dormancy is broken: Supplementary Text Fig. 1.3A-D) in NIL7$^{V79}$ and NIL8$^{A79}$ to the dry-harvest samples from the same PHS trial. We saw a large drop in GE and GI after PHS induction in NIL7$^{V79}$ whereas RGT Planet and NIL8$^{A79}$ showed a less severe germination and PHS phenotype (Supplementary Text Fig. 1.3A-D). This indicates that the hyperactive MKK3$^{V79}$ variant has increased PHS susceptibility (Supplementary Text Fig. 1.4A-D).

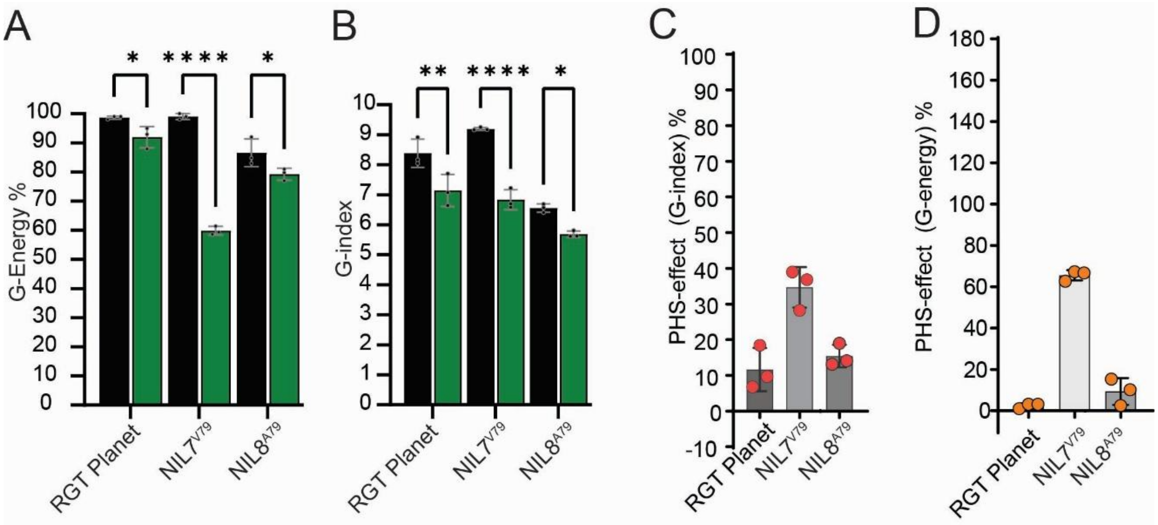

**Supplementary text Fig. 1.3. PHS field trials of Ethiopian *MKK3* haplotype NILs. A,B**, PHS trials of *cv.* RGT Planet (MKK3$^{N383+R350}$) and NIL7$^{V79}$ and NIL8$^{A79}$ in RGT Planet background (PHS field trial, FRA 2023). Shown are (**A**) grain G-energy (GE, in percentage) and (**B**) grain G-index (GI) measured after RGT Planet dormancy is broken. Dry harvest (black bar), and wet harvest (green bar) samples. **C,D**, Calculated PHS-effect (G-energy) in (**C**), and PHS-effect (G-index) in (**D**), of *cv.* RGT Planet, NIL7$^{V79}$ and NIL8$^{A79}$.

Supplementary text 1.5. Wild and Domesticated barley show differences in selection pressure

In wild populations, nonsynonymous substitutions (NSYN), which alter the amino acid sequence of proteins, are typically constrained by purifying selection, as many changes tend to impact fitness negatively and are removed from populations over time. In contrast, silent synonymous substitutions (SYN) are often considered neutral and can accumulate in the genome without impacting fitness (60). In domesticated species, the anthropogenic selection of phenotypically desirable features drives changes in the sequence of genes. The result of positive selection is the accumulation of NSYN that contribute to phenotypic diversification (61, 62). Moreover, domesticated species often suffer from population bottlenecks (reduced population size) leading not only to the fixation of beneficial NSYN variants but also to the increased frequency of mildly deleterious mutations, including SYN, due to genetic drift (61, 62).

We evaluated the impact of domestication on coding sequence diversity at the *MKK3* locus (HORVU.MOREX.r3.5HG0537440) by analyzing SYN, NSYN, and premature stop-codon substitutions per sample in wild and domesticated barley accessions from the PRJEB8044 exome capture dataset.

Here, domesticated barley exhibited a significantly higher proportion of NSYN compared to wild barley (Supplementary text Table S1.5.1, Fig. S1.5.1), with landraces (n = 136) averaging 4.6 NSYN per sample compared to 3.67 in wild accessions (n = 83; p-value = 2.02e-05). This result aligns with the expectation that domestication-induced bottlenecks and directional selection pressures favor environment or end-use-advantageous alleles, such as $MKK3^{T260}$ and $MKK3^{Q165}$, respectively. Conversely, wild accessions showed significantly higher SYN counts (average of 5.17) compared to landraces (average of 3.91; p-value = 3.06e-06), suggesting greater neutral genetic diversity at the *MKK3* locus. Since synonymous mutations do not alter amino acid sequences, they are largely neutral, accumulating primarily through genetic drift, reflecting a less constrained evolutionary history compared to domesticated barley.

**Supplementary text Table 1.5.1. Nucleotide substitutions in wild and domesticated barley accessions.** SYN denotes synonymous substitutions and NSYN nonsynonymous substitutions.

| Substitution type | Group | Sample size | Avg Substitutions | Kruskal Wallis | p_value |
|---|---|---|---|---|---|
| SYN | Domesticated | 136 | 3.91 | 21.78 | 3.06e-06 |
| | Wild | 83 | 5.17 | | |
| NSYN | Domesticated | 136 | 4.6 | 18.17 | 2.02e-05 |
| | Wild | 83 | 3.67 | | |
| STOP | Domesticated | 103 | 1.08 | 1.7 | 0.192 |
| | Wild | 62 | 1.02 | | |

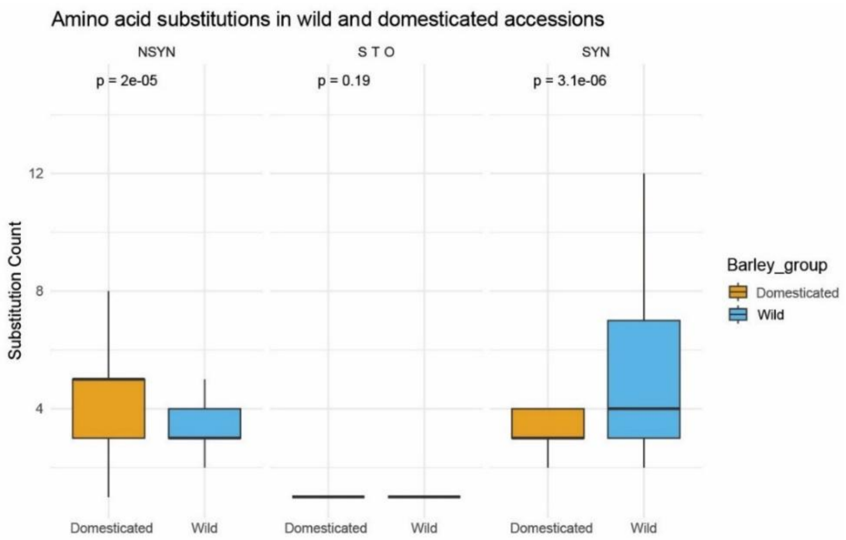

**Supplementary text Figure 1.5.1. Type of nucleotide substitutions and their significance in wild and domesticated barley.** SYN denotes synonymous substitutions, NSYN nonsynonymous substitutions and STO premature-stop substitutions.

The distribution of substitutions between domesticated and wild barley reflects evolutionary dynamics associated with domestication. Domesticated accessions primarily cluster between 3 and 5 NSYN per sample, with some examples reaching some 10–12 substitutions per sample. Wild barley accessions tend to have 3–5 NSYN but exhibit fewer extreme values, indicating stronger purifying selection pressures that limit the accumulation of deleterious variants. In terms of SYN, domesticates peak between 2–4 substitutions per sample, indicative of reduced neutral genetic diversity due to population bottlenecks. In contrast, wild barley displays a broader distribution (2–8 SYN), reflecting greater neutral diversity maintained by larger effective population sizes and less intense selection pressure. Overall, these patterns suggest that domestication events have shaped genetic diversity by promoting the fixation of beneficial nonsynonymous mutations while reducing synonymous diversity through genetic drift and bottlenecks.

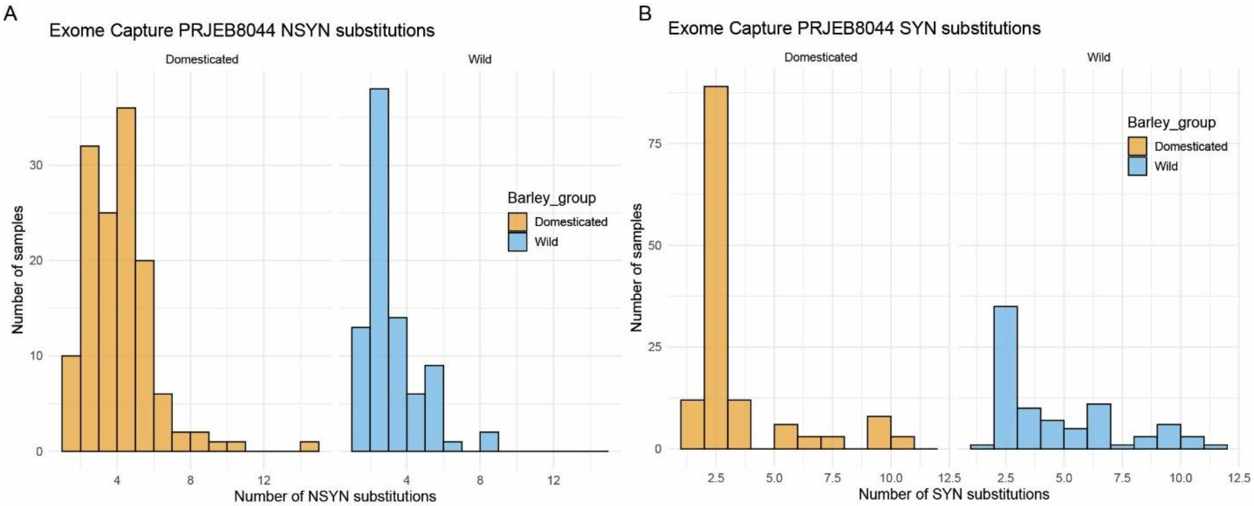

**Supplementary text Fig. 1.5.2. Frequency of nucleotide substitutions in wild and domesticated barley. A,B**, distribution of (A) NSYN and (B) SYN substitutions across the exome captured (PRJEB8044) wild and domesticated barley diversity panel.

We further investigated the distribution and domain-location of identified amino acid variants in the MKK3 peptide sequence. The MKK3 protein model suggests several functional domains and regions likely crucial for enzyme activity and protein-protein interactions (Fig. 3D). The intrinsically disordered protein region (20–39 aa) may be involved in flexible interactions and signaling regulation. The kinase domain (89–343 aa) is essential for catalytic activity and enzymatic regulation, and the NTF2 Domain (372–522 aa) is potentially involved in nuclear transport or additional regulatory functions. Domesticated barley exhibits amino acid variants across all three protein domains, consistent with the expectation of relaxed constraint or positive selection during domestication (Supplementary text Figure 1.5.3). By contrast, the reduced number of amino acid variants in the kinase domain among wild barley accessions likely reflects purifying selection acting on conserved functional domains of MKK3. This pattern aligns with genomic signatures of selection described in domestication genes such as *tga1*, *sh4*, and *Q* in other cereals (61), where protein-level changes in key regulators facilitated phenotypic transitions. Overall, this highlights how domestication has targeted specific functional regions of MKK3, promoting beneficial mutations while preserving essential structural integrity necessary for protein functionality.

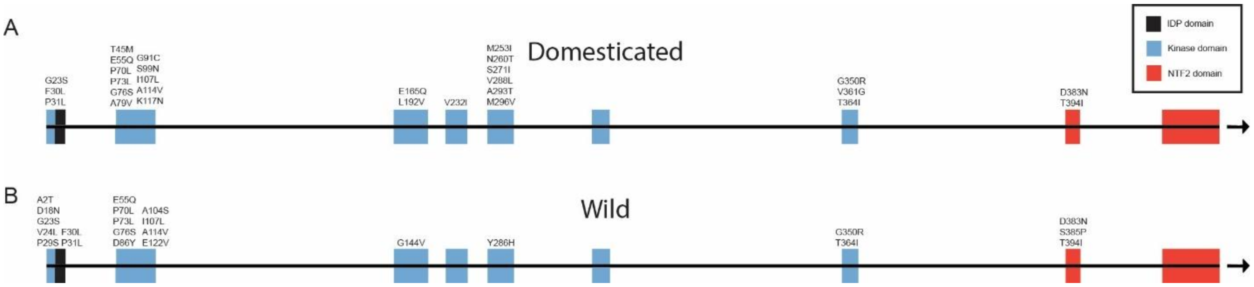

**Supplementary text Figure 1.5.3. Amino acid variants in MKK3. A,B,** Distribution of amino acid variants across the MKK3 peptide sequence in (**A**) domesticated and (**B**) wild barley accessions. Predicted protein domains are colored in black (IDP domain), blue (kinase domain) or red (NTF2 domain). Notably, key domestication-related nonsynonymous substitutions, such as N260T and E165Q occur within the kinase domain, critical for MKK3 function.

### Figs. S1 to S7

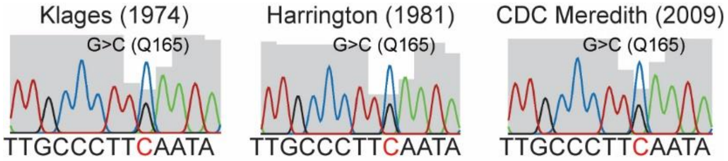

**Supplementary Fig. S1. Genotyping *MKK3* in North American barley**. Sanger sequencing results from nucleotide position 485-497 with a heterozygous chromatogram signal at position 493 (numbering relative to $MKK3^{Ref}$ reference sequence HORVU.MOREX.PROJ.5HG00474740, see Supplementary Table S2; grey shading denotes sequence quality) in barley *cvs.* Klages, Harrington and CDC Meredith (shown with respective year of variety registration).

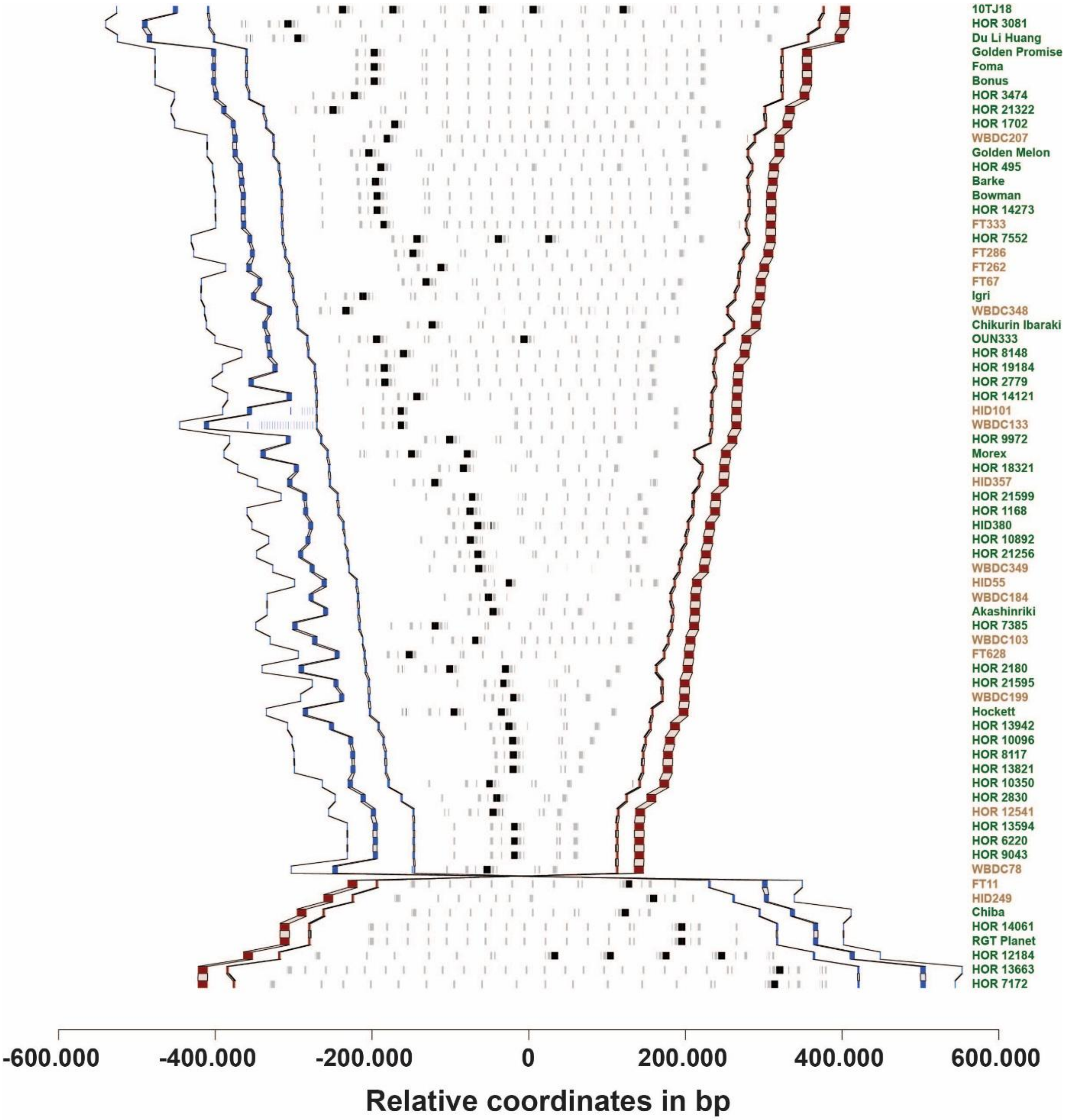

**Supplementary Fig. S2. *MKK3* structural variation in 76 BPGv2 genome assemblies.** *MKK3* in black, other genes in grey. Red and blue squares denote marker genes that define the synteny, delimit the region and sort the accessions based on the distance between endpoints. Lines connect gene models between different genomes. Accession names are given on the right axis. One accession (HOR 3365) has a very large insertion (not shown). In four accessions (FT11, Chiba, RGT Planet and HOR 3365), the larger MKK3 region is inverted (6, 4, 4 and 2.5 Mb, respectively). Barley accessions are coloured according to their domestication status with wild in light brown and domesticated in green.

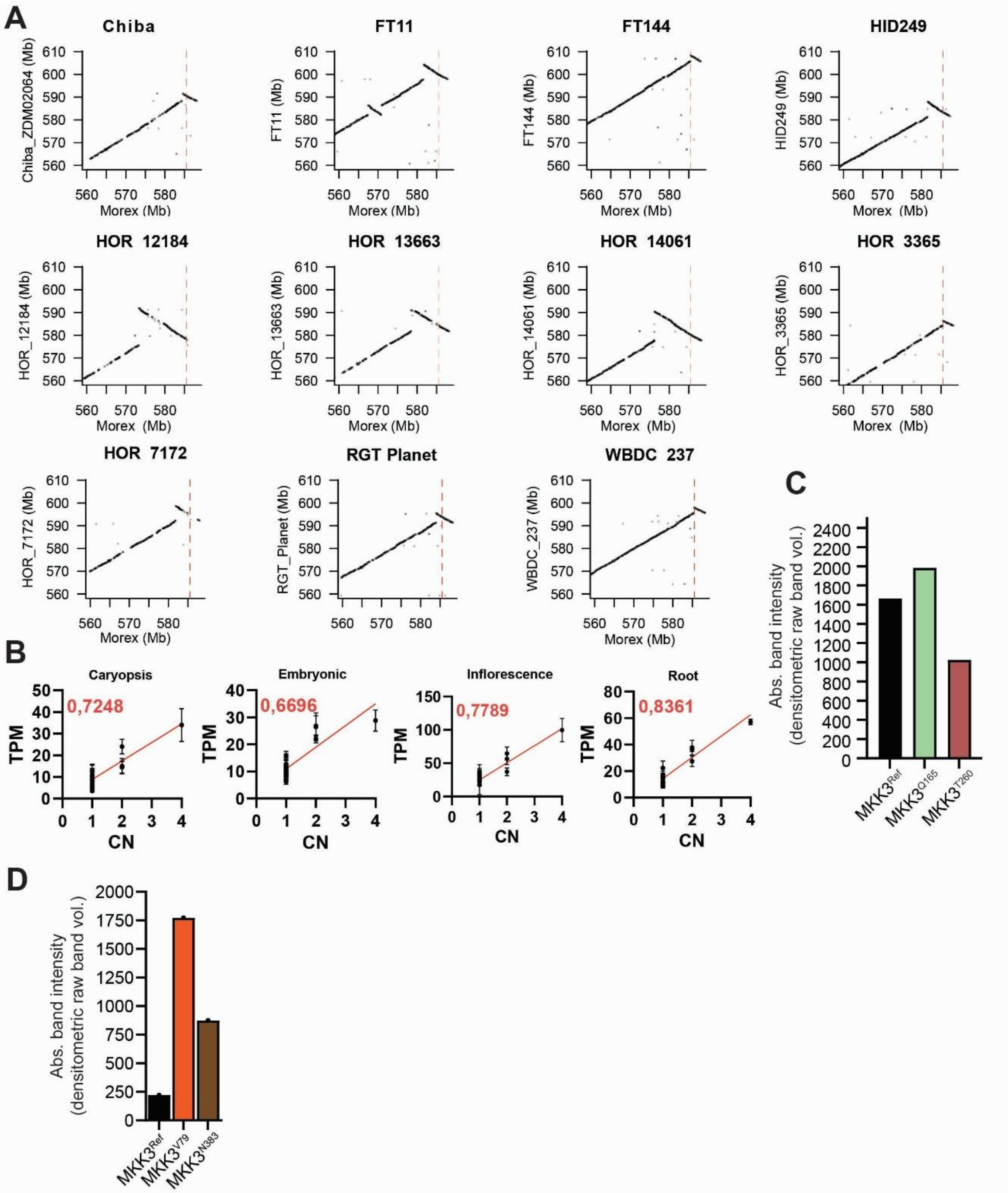

**Supplementary Fig. S3.** ***MKK3*** **gene analysis, kinase activities and origin of accessions with high CN. A**, Dot plot alignments of the *MKK3* genome region of chromosome 5HL from BPGv2 genome assemblies with *MKK3* chromosomal inversions against *cv.* Morex. **B**, *MKK3* transcript abundance in the pan-transcriptome (*29*) in caryopsis, embryonic, inflorescence, root, and total-tissue as a function of copy number with $r^2$ value of the linear regression fit in red. **C**, *In vitro* kinase activities of $MKK3^{Ref}$ (Ref sequence, HORVU.MOREX.PROJ.5HG00474740; Supplementary Table S2), $MKK3^{T260}$ and the $MKK3^{Q165}$ variants (n=1). **D**, *In vitro* kinase activity of $MKK3^{Ref}$ (Ref sequence, HORVU.MOREX.PROJ.5HG00474740; Supplementary Table S2), $MKK3^{V79}$ and the $MKK3^{N383}$ variants (n=1).

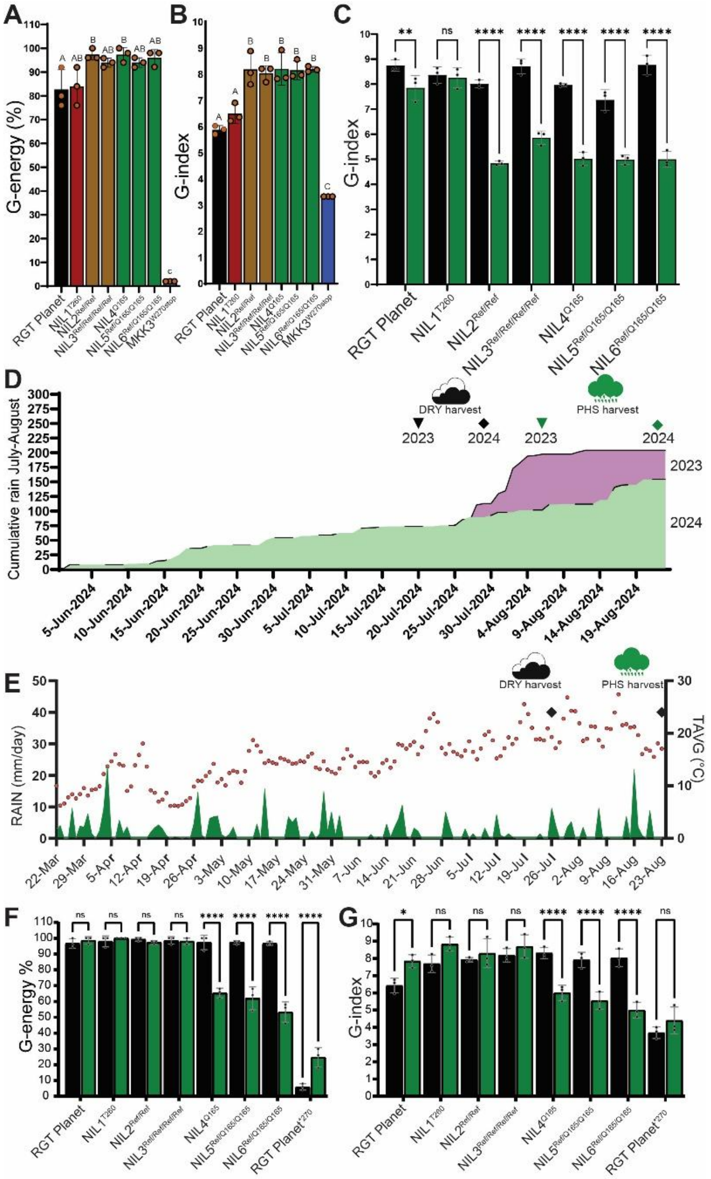

**Supplementary Fig. S4. Extended PHS field trials and analyses of diverse *MKK3* haplotype NILs**. **A,B**, Measurements of germination energy **(A)** and germination index **(B)** to assess dormancy at week 3 after harvest of *cv.* RGT Planet (MKK3$^{R350+N383}$), NIL1$^{T260}$, NIL2$^{Ref/Ref}$, NIL3$^{Ref/Ref/Ref/Ref}$, NIL4$^{Q165}$, NIL5 $^{Ref/Q165/Q165}$, NIL6$^{Ref/Q165/Q165}$ (field grown, FRA 2024). **C**, PHS trials of *cv.* RGT Planet, NIL1$^{N260T}$, NIL2$^{Ref/Ref}$, NIL3$^{Ref/Ref/Ref/Ref}$, NIL4$^{Q165}$, NIL5$^{Ref/Q165/Q165}$, NIL6$^{Ref/Q165/Q165}$ (field grown, FRA 2023). G-index is shown after *cv.* RGT Planet dormancy is broken. Dry harvest (black bars), and wet harvest (green bars) samples. **D**, Cumulative rain (mm) in the months of June, Juli and August across the years 2023 (lilac) and 2024 (green) shading. **E**, Weather data [the average air temperature (TAVC) in degree Celsius in red circles and precipitation (RAIN) in mm/day in green shading] for PHS field trial site in FRA in the growth season 2024. Marked with squares are the harvest time points of 'dry' (26$^{th}$ of July) and 'PHS' samples (23$^{rd}$ of August). **F-G**, PHS trials of *cv.* RGT Planet (MKK3$^{R350+N383}$), NIL1$^{T260}$, NIL2$^{Ref/Ref}$, NIL3 $^{Ref/Ref/Ref/Ref}$, NIL4 $^{Q165}$, NIL5 $^{Ref/Q165/Q165}$, NIL6 $^{Ref/Q165/Q165}$ , RGT Planet$^{270*}$ (field-grown, FRA 2024). Shown are grain A) G-energy (percentage) and B) G-index after *cv.* RGT Planet dormancy is broken. Dry harvest (black bar) and PHS harvest (green bar) (*$P \leq 0.05$, ** $P \leq 0.01$, *** $P \leq 0.001$, and **** $P \leq 0.0001$; ns, not statistically significant).

Placeholder
Figure attached as PDF

**Supplementary Fig. S5. Median-joining haplotype network of *MKK3* copies in 228 exome sequenced accessions (83 wild and 145 domesticated) (56).** Nodes representing haplotypes found in one or multiple accessions are labelled with the accession name or a # in bold (Supplementary Table S8), respectively, followed by the amino acid variation relative to $MKK3^{Ref}$ (Ref sequence, HORVU.MOREX.PROJ.5HG00474740) (28). The node size is proportional to the number of gene IDs a given node represents.

Placeholder
Figure attached as PDF

**Supplementary Fig. S6. Clustering analysis of the larger *MKK3* region using 50K-SNP-array data.** Three Bere barley containing subclades are highlighted in green, blue and red.

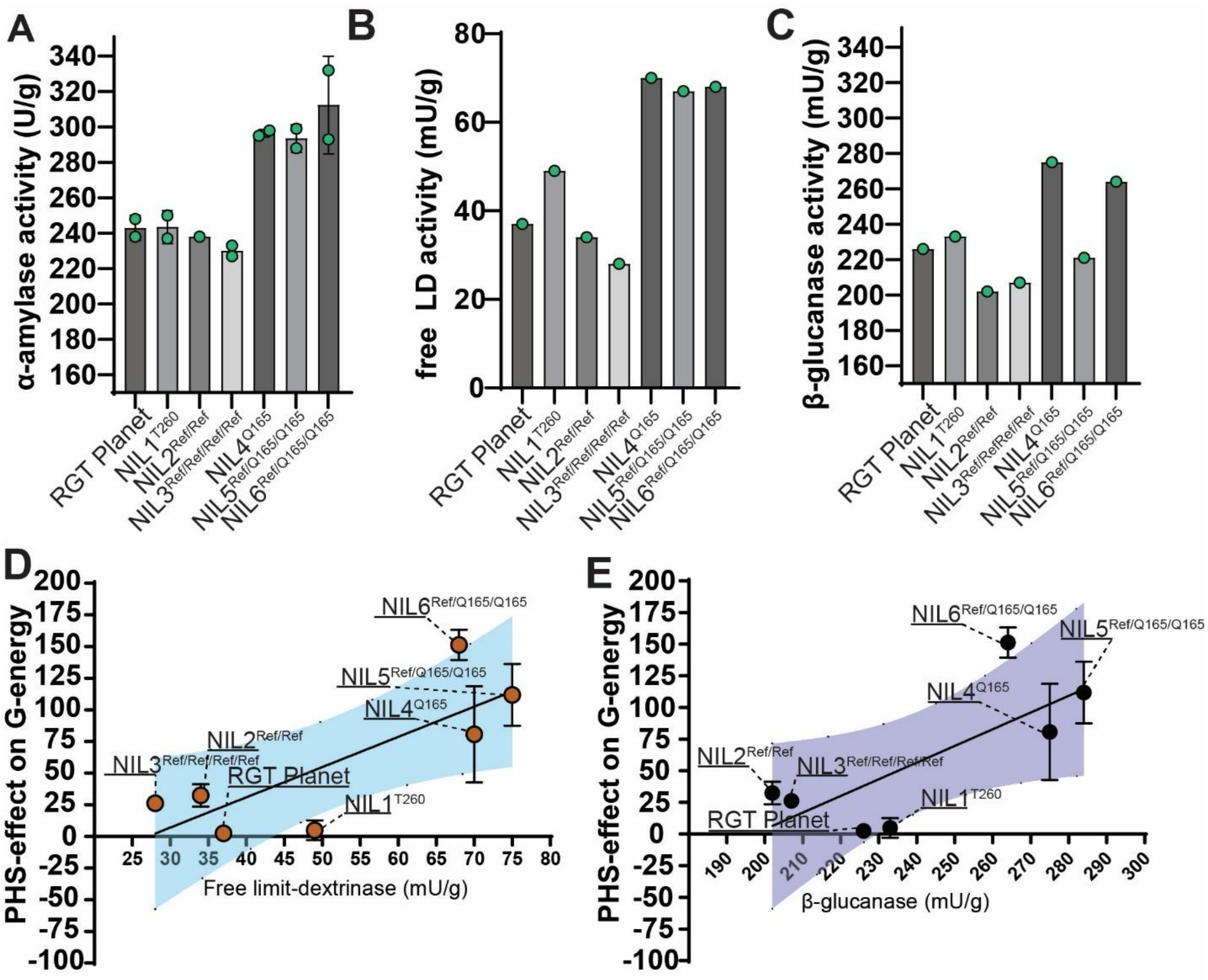

**Supplementary Fig. S7. Micro-malt analyses of *MKK3* NILs and RGT Planet control . A**, α-amylase activity, **B**, free limit dextrinase (LD) activity and, **C**, β-glucanase activity (n=1). Plants were field grown in NZL 2023. **D-E**, PHS-effect (G-energy, calculated from PHS data shown in Fig. 4D, see Materials) as a function of **(D)** free limit dextrinase activity or (**E**) β-glucanase activity of micro malted field grown grains (NZL 2024).

**Supplementary Table S1.** ddPCR-based *MKK3* copy number (CN) analysis and genotyping results for variants $MKK3^{V79}$, $MKK3^{E165}$, $MKK3^{Q165}$, $MKK3^{T260}$. Entries are grouped by topic and some entries may be duplicated.

**Supplementary Table S2.** *MKK3* haplotype analysis in BPGv2 (28).

**Supplementary Table S3.** *MKK3* copy number analysis and genotyping across a barley diversity panel (28) using k-mer and ddPCR approaches. See row 1712 for column description.

**Supplementary Table S4.** *MKK3* genotyping matrix for Exome Capture PRJEB8044 (56). Presence/absence of non-synonymous SNPs across accessions based on k-mer-specific genotyping.

**Supplementary Table S5.** *MKK3* GsRTDs and transcripts linked to BPGv2 gene IDs, amino acid haplotypes and 5HL inversions from the pan-transcriptome (29).

**Supplementary Table S6.** ddPCR based *MKK3* genotype analysis of NIL donor accessions. *MKK3* copy number analysis using ddPCR and sequence analysis of sequenced MKK3 haplotypes (56).

**Supplementary Table S7.** *MKK3* copy number analysis across a barley diversity panel (45) using k-mer and ddPCR approaches.

**Supplementary Table S8.** Haplotype network legend to Supplementary Fig. S5. Accession names associated with node #.
