## Supplementary figures and images for "Post-Domestication selection of MKK3 Shaped Seed Dormancy and End-Use Traits in Barley"

### Supplementary Fig. S5

#### Supplementary Fig. S5

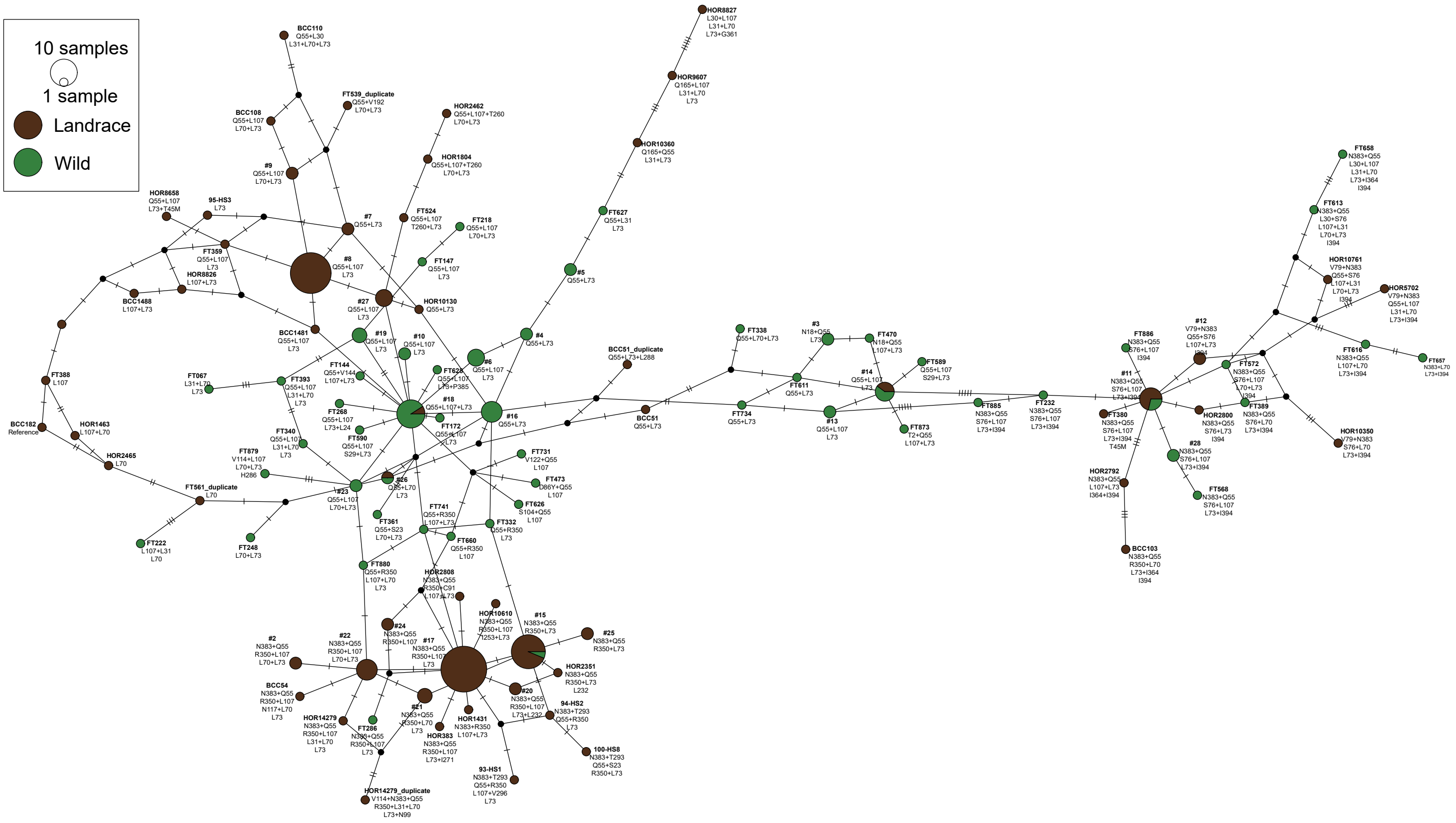

### Supplementary Fig. S6

Supplementary Fig. S6

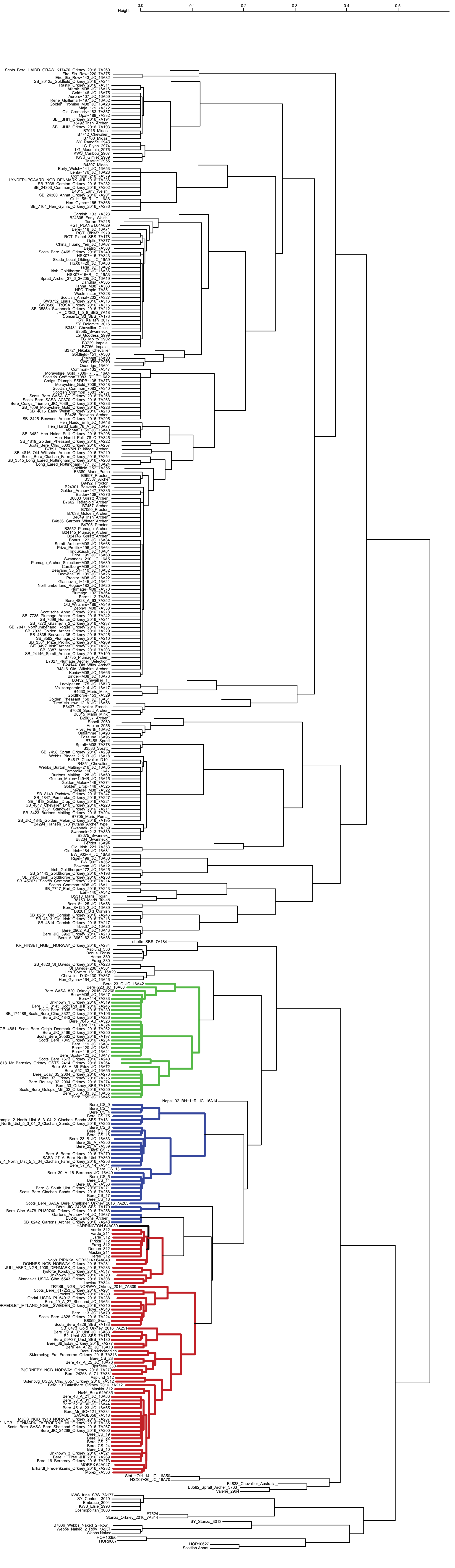
